# A geometric approach to scaling individual distributions to macroecological patterns

**DOI:** 10.1101/302810

**Authors:** Nao Takashina, Buntarou Kusumoto, Yasuhiro Kubota, Evan P. Economo

## Abstract

Understanding macroecological patterns across scales is a central goal of ecology and a key need for conservation biology. Much research has focused on quantifying and understanding macroecological patterns such as the species-area relationship (SAR), the endemic-area relationship (EAR) and relative species abundance curve (RSA). Understanding how these aggregate patterns emerge from underlying spatial pattern at individual level, and how they relate to each other, has both basic and applied relevance. To address this challenge, we develop a novel spatially explicit geometric framework to understand multiple macroecological patterns, including the SAR, EAR, RSA, and their relationships, using theory of point processes. The geometric approach provides a theoretical framework to derive SAR, EAR, and RSA from species range distributions and the pattern of individual distribution patterns therein. From this model, various well-documented macroecological patterns are recovered, including the tri-phasic SAR on a log-log plot with its asymptotic slope, and various RSAs (e.g., Fisher ⊠ s logseries and the Poisson lognormal distribution). Moreover, this approach can provide new insights such as a single equation describing the RSA at an arbitrary spatial scale, and explicit forms of the EAR with its asymptotic slope. The theory, which links spatial distributions of individuals and species with macroecological patterns, is ambiguous with regards to the mechanism(s) responsible for the statistical properties of individual distributions and species range sizes. However, our approach can be connected to mechanistic models that make such predictions about lower-level patterns and be used to scale them up to aggregate patterns, and therefore is applicable to many ecological questions. We demonstrate an application of the geometric model to scaling issue of beta diversity.

## 1 Introduction

The problem of pattern and scale is central in ecology, and critical insights into conservation biology emerge as we observe different aspects of ecosystems across scales [1]. The species area relationship (SAR), endemic area relationship (EAR), and relative species abundance (RSA) are such examples characterizing macroecological and community patterns of ecosystem across scales. Each of these patterns has a long history and ample literatures (e.g., [2–6]), and there are still active discussions over fundamental patterns (e.g., [7, 8]). In particular, recent researches have focused on the quantitative investigation of scaling issues of macroecological patterns [9-12]. However, little is known about how individual-level spatial distributions scale up to aggregate patterns such as the SAR, EAR, and RSA (but see [11]).

A model linking individual distributions with aggregate patterns can provide a great applicability to other ecological questions in community ecology and conservation biology. For example, to investigate the scale-dependence of beta diversity [13], one needs species abundance information at finer scales and a consistent scale-change operation must be defined. These macroecological patterns are also critical for implementing effective ecosystem management, given accelerating loss of biodiversity worldwide [14, 15]. For example, the SAR and EAR provide information about how many species will be affected and be lost from the landscape when a certain region is degraded, respectively [16], and the RSA provides more detailed information on community structure such as composition of rare species. Practically, conservation decision making is spatially explicit provided optimal implementation strategy of protected areas with budget constraints [17, 18], and often high degree of spatial information, such as individual distributions and the number of individuals across multiple scales are required (e.g., [19, 20]).

Here, we propose a novel geometric approach to study macroecology that provides a bridge between spatial distribution patterns and emergent macroecological patterns. In this framework, the SAR, EAR, and RSA are derived from the geometry of species ranges and the spatial distribution of conspecific individuals therein. Conceptually similar approaches have been investigated in previous researches without individual distributions [9, 10, 21], and with explicitly integrating individual distributions [11, 22]. Allen and White [21] derived an upper bound of the SAR, by calculating an overlapping area between the sampling region and the randomly placed distribution range without considering individual distributions therein, leading to a biphasic SAR on a log-log plot. Although this model provides the well-known asymptotic slope of 1 at very large sampling scales, it did not capture the sampling process on smaller scales responsible for the triphasic curve. Plotkin et al. [22] applied the Thomas process, a point process model, to generate aggregated individual distribution patterns in tropical forest plots. They independently estimated a dispersal distance and density from spatial distribution of individual trees of each species, and generated superposition of multiple species in a forest. The fitted Thomas process model very precisely recovers SARs up to 50- ha. Grilli et al. [11] also applied a point process model to account for individual distributions. With aggregated communities generated by the Poisson cluster model, they derived the SAR, EAR, and RSA. In doing so, the model requires a neutral assumption and a specific input of the RSA form in the whole system.

In our framework, the model uses a point field of communities that holds information of all individuals and species distributions. The point field is consistent with scale changes and, therefore, we can derive macroecological patterns across scales consistently without changing ecological parameters used. This property also provides a significant applicability to conservation practices, since the model provides macroecological information on arbitrary spatial scales. Particularly, we assume non-interacting species distribution ranges and individual distributions therein. In derivations of the SAR, EAR, and RSA, our model put a particular emphasis on geometric patterns that the sampling region, distribution ranges, and its individual distribution patterns therein generate. One of the advantages of this approach is that we can discuss in what conditions the model recovers emergent macroecological patterns, and, importantly, any preceding knowledge of these patterns are required unlike the previously developed model [11]. The model is also amenable to incorporate variations in species characteristics, and we can easily discuss the emergent patterns when species vary in ecological characteristics.

The paper contains three major parts. First, we develop a general theory based on our geometric approach to link the SAR, EAR, and RSA to spatial point patterns, without assuming specific spatial patterns. This provides a rather general framework, and any spatial patterns define species and individual distribution pattern can be applied. Second, we derive emergent macroecological patterns by assuming specific point patterns that phenomenologically describe geometric patterns of species and individual distributions therein. We show the model recovers many well-documented macroecological patterns including the tri-phasic SAR with its asymptotic slope 1 on a log-log plot. For RSAs, well-known RSAs such as negative binomial distribution, Fisher’s logseries, and the Poisson lognormal are recovered at the small sampling scales. We demonstrate that at intermediate sampling scales, RSAs are described by combinations of two probability distribution functions, and it leads to a left-tailed or negatively-skewed bell shape curves on a logarithmic scale. Notably, these all RSAs are summarized by a single equation, and therefore easy to discuss transition patterns with sampling size. It is worth noting that the structure of these SAR and RSA patterns are often attributed to ecological and evolutionary mechanisms, but our geometric approach still can recover these patterns based on general assumptions about individual and range size distributions. Our model also provides new potential form of EARs with asymptotic slope 1 on a log-log plot that, unlike commonly used definition, directly counts the average number of enclosed species distributions. Third, to demonstrate potential applicability of the geometric model to ecological questions, we address the scale-dependence of beta diversity as an example.

## 2 General theory

We develop here a general theory of our geometric approach, an attempt to provide a general framework for calculations of the SAR, EAR, and RSA without assuming any specific geometric properties. More specifically, here we do not assume individual distribution patterns,
shapes of the sampling region and the distribution range. However, it is worth noting that the central assumptions of the theory are that we assume no interaction between species and, therefore, species distribution range of each species are chosen independently in the homogeneous environment. Once ecological properties and the sampling scheme are specified, the SAR, EAR, and RSA can be obtained via the general framework. We will discuss some specific situations in Results. Central parameters used throughout the paper are summarized in Table 1.

**Table 1:** Definition of parameters

| Symbol | Parameter |
| --- | --- |
| $S$ | Sampling region |
| $D, (D_i)$ | Distribution range of species ($i$) |
| $\nu(A)$ | Area of $A$ |
| $\mathbf{x}_c^A$ | Geometric center of $A$ |
| $l_A$ | Minimum distance between the center and boundary of $A$ |
| $L_A$ | Maximum distance between the center and boundary of $A$ |
| $X$ | Number of individuals |
| $N_i(A)$ | Number of individuals of species $i$ in $A$ |
| $N_s(A)$ | Number of species in $A$ |
| $\lambda_i$ | Intensity of species $i$ |
| $\lambda_s$ | Intensity of species |
| $\mu_i(A)$ | Intensity measure of species $i$ in $A$ |
| $\mu_s(A)$ | Intensity measure of species in $A$ |

### 2.1 Basic notation

#### 2.1.1 Intensity and intensity measure

We introduce some basic quantities of ecosystem which will be used in the following discussions. Each of these are generally used in the field of point process, and we borrow some ideas in this paper. Interested readers may refer to literatures in the field (e.g., [23, 24]) or applications to ecological studies (e.g., [11, 19, 20, 22, 25]). Later, we will combine the general theory with point processes to examine specific individual distribution patterns such as random or clustering distributions.

The intensity, λ*_i_*, and the intensity measure, *μ_i_*, of species *i* are the central quantities of our model characterizing the ecological community: the average density of individuals of species *i*, and the average number of individuals of species *i* in region *A*, respectively. Given the area *ν*(*A*) and the number of individuals of species *i*, *N_i_*(*A*) in *A*, the intensity of species *i* in *A* is defined as [24]

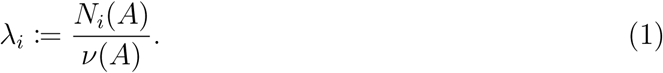

The intensity measure of species in *A* is given by multiplying the intensity by area *ν*(*A*)

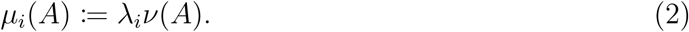

The same quantities for species assembly are also defined within this framework. Let *N_s_*(*A*) be the number of species in *A*. Then, the intensity and intensity measure in *A* of the species assembly are defined as λ*_s_* := *N_s_*(*A*)/*ν*(*A*) and *μ_s_* := λ_*s*_*ν*(*A*), respectively. Then the vector **λ** = (λ_1_ λ_2_ … λ_*s*_) holds the information of community abundance. Let us define the intensity measure of species, *μ_s_*(*A*) as

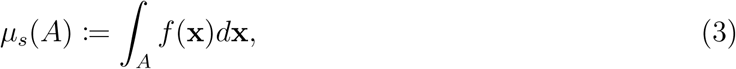

where, *f* (x) is a point field holding information on probability of all individual locations in two-dimensional space x ∈ **R**^2^. Let us assume the homogeneous environment, and then we can put *f* (x) = λ_*s*_, where λ_*s*_ is the intensity of species, and we obtain

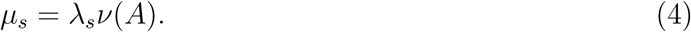

#### 2.1.2 Sampling region and endemic region

In this paper, we refer to *S* as the *sampling region* which defines the spatial scales discussed. Similarly, we refer to *D_i_* as the distribution range of species *i.* We often omit the subscript *i* and call the *distribution range of species* or simply *distribution range* if obvious. Throughout the analysis, the distribution range *D* is used as an endemic region of species where the individual distributions of the species are restricted within the region. For convenience, yet mathematically not rigorous, we use the symbols “ > “ and “ < ” regarding to relative sizes of *S* and *D*. For example, in our notations *S* > *D* implies that we can always find a shift, without rotating, that satisfies *D* ⊂ *S*.

### 2.2 Species area relationship

Here we derive a method to calculate the species area relationship (SAR). Often it exhibits the tri-phasic features on a log-log plot [3, 5], showing a fast increase on small scales, slowing down the increase on intermediate scales, and again shows a quick increase asymptotically toward to 1 for the slope on very large scales where sampling region exceeds correlation distance of biogeographic process with separate evolutionary histories [5, 6]. As the simplest example, let us first assume a neutral community where biological properties, such as intensity of each species, distribution range, and average dispersal distance are identical between species. However, as we will see below, it is straightforward to incorporate species variations. To calculate the species number in a given area, we use the nonzero probability 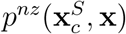 that a species at x with the geometric center of the distribution region 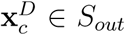 provides at least one individual to the sampling region *S* with its geometric center 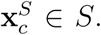 *S_out_* is the region that if the geometric center of the distribution range of a species *D* belongs to this region, 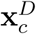 ∈ *S_out_*, the species potentially provides its individuals to the sampling region *S* (Fig. 1). The sampling region *S* is the subset of *S_out_*, *S* ⊂ *S_out_*. In this simplest example, the average number of species given sampling area *ν*(*S*) is described:

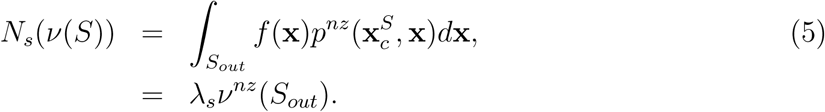

**Figure 1:**
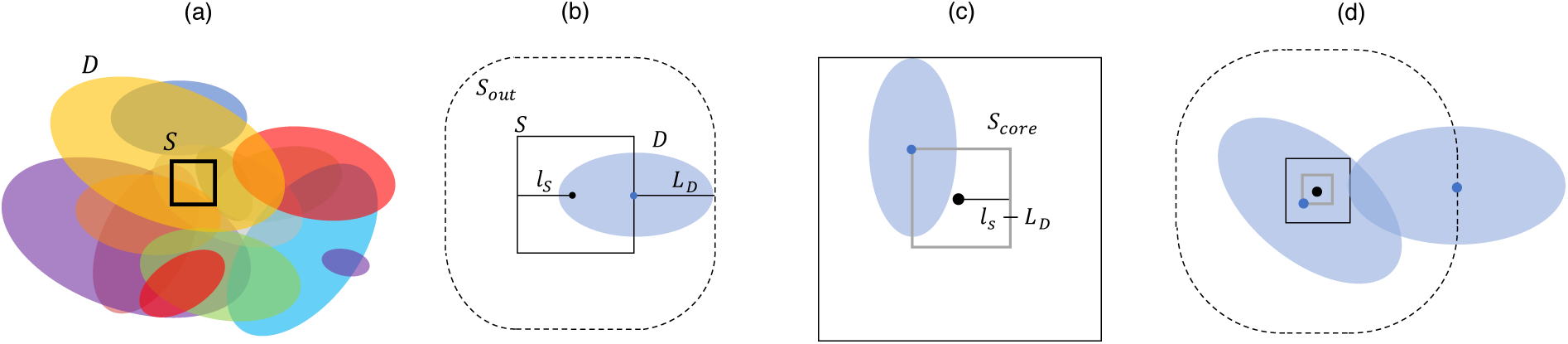
Schematic representations of (a) the sampling region and distribution ranges, and typical geometric relationships used in derivations of the (b) SAR, (c) EAR, and (d) RSA. For each panel, *S* and *D* are the sampling region and distribution ranges, respectively. *S* and *D* have arbitrary shapes in reality, but they are shown by square (black) and ellipse (filled by color) with the geometric center, respectively as an example. *S_out_* is the region described with dotted line, where if the geometric center of a distribution range 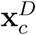 falls in this region, then the probability to overlap *S* and *D* is not zero: Namely *P*(*S* ∩ *D*) ≠ 0 provided 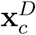 ∈ *S_out_*. *S_core_* is the region described by grey line where provided *S* > *D* (*S* < *D*) if the distribution center of a species 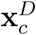 falls in this regions, then the entire distribution range *D* (*S*) is covered by *S* (*D*): Namely *P*(*D* ⊂ *S*) = 1 (*P*(*S* ⊂ *D*) = 1) provided 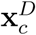 ∈ *S_core_*. For the derivation of the EAR, only the case *S* > *D* is used, but both *S* > *D* and *S* < *D* are used for the derivation of the RSA. *S_core_* is the closed region and typically a subset of the sampling region *S_core_* ⊂ *S*, in which any points on this boundary, x ∈ *∂S_core_* holds the following relationship: when *S* > *D*, min{∥x – x′∥;x ∈ *∂S*, x′ ∈ *∂S_core_*} = max{∥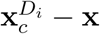∥; x ∈ *∂D*} when *S* < *D*, min{∥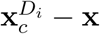∥; x ∈ *∂D*} = max{∥x – x′∥;x ∈ *∂S*, x′ ∈ *∂S_core_*}. Each region is determined by the minimum distance *ℓ_S_*, *ℓ_D_* and maximum distance *L_S_*, *L_D_* between the sampling center and its boundary. See further explanations in main text.

Since the first differentiation of area is proportional to length, the final output of Eq. (5) represents the area of region *S_out_* weighted by nonzero probability *p^nz^*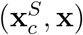 over the region *S_out_*. The SAR is obtained by calculating Eq. (5) across sampling areas *ν*(*S*). From Eq. (5), we can show that the slope of the SAR on a log-log plot approaches 1 at large scales. Firstly, the slope is calculated by

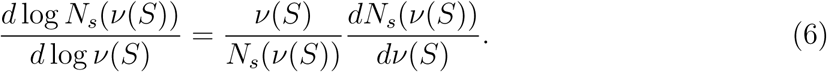

Secondly, when the sampling region becomes significantly larger than the distribution range *S* ≫ *D*, *S_out_* is nearly equivalent to *S*. In this limit, *p^nz^*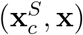 ≃ 1 is satisfied, providing

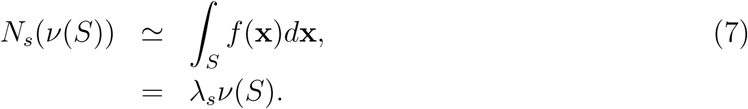

The slope 1 is obtained by substituting this into Eq. (6). Due to the prerequisite relationship *S* ≫ *D*, this asymptotic slope is achieved faster if the distribution ranges are smaller.

As noted above, it is easy to incorporate species variations by assuming that some biological properties follow certain probability distribution functions (pdfs). It is suffice to consider biological parameters that affect the non-zero probability *p^nz^* 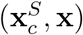 (e.g., distribution range and intensity of each species). We define such parameters by a vector b = (*b*_1_,*b*_2_, …,*b_k_*) ∈ *B*, where *B* is a biologically plausible parameter space. Here, let us assume the independence of biological parameters **b** from the distribution ranges *D*. *D* varies between species, and hence it generates a probability distribution of *S_out_*, *p*(*S_out_*). Therefore, the probability distribution functions of biological parameters and the distribution range are described by *p*(**b**, *S_out_*) = *p*(**b**)*p*(*S_out_*). Now, Eq. (5) becomes

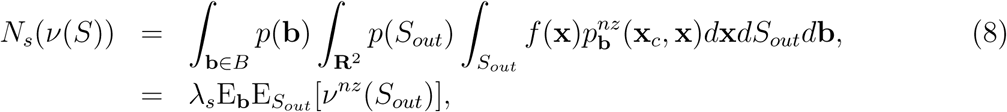

where, E*_S_out__* and E_**b**_ are the average over the distribution ranges and biological parameters, respectively. As *ν^nz^*(*S_out_*) is the average area to find at least one individual, E_**b**_E*_S_out__* [*ν^nz^* (*S_out_*)] is the such area averaged over the pdfs *p*(**b**) and *p*(*S_out_*). Eq. (5) is simply a special case of Eq. (8) by setting *p*(**b**) = *p*(*S_out_*) = 1 with choosing a set of parameters. We can make intuitive discussions about the effect of biological parameters from Eq. (8). For example, if the intensity of each species is high, it is likely to increase the area *ν^nz^*(*S_out_*) and it increases the number of species in given area, and vice versa. In this situation, the same discussion about the slope of SAR curve Eq. (7) can be made as long as the sampling region is significantly larger than distribution range of each species, *S* ≫ *D_i_* for all *i*.

### 2.3 Endemic area relationship

Next we discuss the derivation of the endemic area relationship (EAR). Despite its importance to conservation biology, understanding of the empirical shape across scales is largely deficient [10]. Here, we define the endemic species, with explicitly introducing distribution region with an arbitrary shape, as the species with its distribution range *D* is covered by the sampling region *D* ⊂ *S*. This definition is different from the commonly used definition (e.g., [10, 11, 16]) where the endemic species is defined, using the SAR, as the average number of species these population is completely covered by area *A*: *N_en_*(*A*) = *N_s_*(*A*′) – *N_s_*(*A*′ \ *A*), where *N_en_*(*A*) is the number of endemic species in area *A*, and *A*′ is the system size. This definition relies on the occurrence probability of individuals of a species, and hence its individual distribution patterns. Typically SAR curves show *N_s_*(*A*) > 0 with *A* > 0, and therefore *N_en_*(*A*) > 0 for any small *A*. On the other hand, our definition is based on an explicit form of the endemic region, and the number of the endemic species is 0 if the sampling region is smaller than distribution range of all species: *S* < *D_i_* for all *i.* In this framework, individual distribution patterns do not affect the EAR. This definition may be more straightforward to discuss an extinction of species, since habitable regions of all species are explicitly taken into account. However, as we will see below, the two definitions become equivalent in the large limit of *A*.

In our framework, the number of endemic species given area is calculated as follows

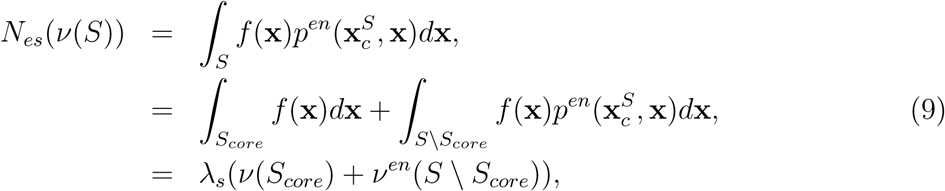

where, *p^en^*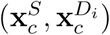 is the endemic probability that a distribution range with its geometric center 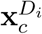 ∈ *S* \ *S_core_* becomes subset of the sampling region with the geometric center 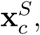 *D_i_* ⊂ *S*. *S_core_* is the subset of the sampling region *S* where if the distribution center of a species 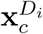 falls in this regions, then the entire distribution range *D_i_* is covered by *S*: *P*(*D_i_* ⊂ *S*) = 1 provided that 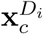 ∈ *S_core_* (Fig. 1). As in the case of the SAR, we can show the slope of the EAR on a log-log plot approach 1 when *S* ≫ *D.* In this limit, *S_core_* is nearly equivalent to *S*, and hence Eq. (9) becomes

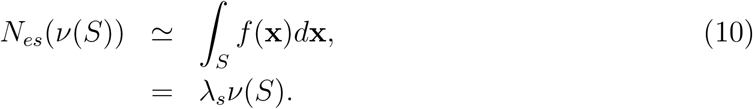

This is equivalent to Eq. (7), and hence the slope approaches 1. As we noted, the same result can be obtained in another definitions using the SAR, since *N_en_*(*A*) ≃ *N_s_*(*A*) when *A* approaches *A*′.

We now introduce the EAR in the situation where species have different distribution areas *ν*(*D_i_*). This generates a probability distribution of the core region of sampling, *p*(*S_core_*). By introducing the variation in distribution areas, Eq. (9) now becomes

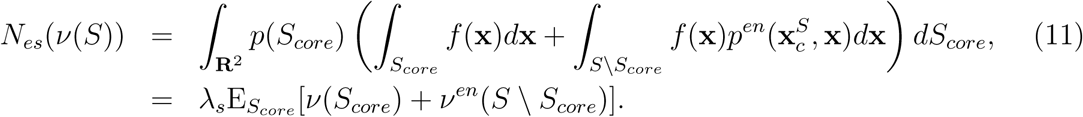

As in the case of the SAR, we obtain Eq. (9) by setting *p*(*S_core_*) = 1 in Eq. (11). In addition, the same discussion about the slope of the EAR can be made in this situation, if the sampling region is significantly larger than the distribution ranges: *S* ≫ *D_i_* for all *i*.

### 2.4 Relative species abundance

Here, we develop a method to derive the relative species abundance (RSA). RSA, or SAD, a histogram of species count, is a product of multiple ecological mechanisms such as species interactions and stochasticity [7]. For convenience, we use the notation *P*(*X* = *x*) to describe the probability of finding a species with abundance *x.* Derivation of the RSA is more cumbersome than the SAR and EAR since we use all probabilities of *P*(*X* = *x*). In our framework, the average abundance of species *i* is proportional to the area where the sampling region *S* and the distribution range of the species *D_i_* are overlapped, *ν*(*S*∩*D_i_*). Each sampled species has different overlapped area (*ν*(*S* ∩ *D_i_*) ≠ *ν*(*S* ∩ *D_j_*), (*i* ≠ *j*)), and its area changes as sampling and distribution scales as well as these shapes change. Therefore, to calculate the RSA across scales we need a pdf of *ν*(*S* ∩ *D*), provided *ν*(*S* ∩ *D*) ≠ *ϕ*, which describes the variation of *ν*(*S* ∩ *D*) between sampled species.

Roughly speaking, there exist two regimes to determine the maximum value of *ν*(*S* ∩ *D*) among sampled species: *S* > *D* and *S* < *D*, provided *ℓ_s_* > *L_d_* and *L_s_* < *ℓ_d_*, respectively. Namely, the maximum overlapped area is either *ν*(*S*) or *ν*(*D*) depending on *S* < *D* (Fig. 2a, b) or *S* > *D* (Fig. 2c, d), respectively: max{*ν*(*S* ∩ *D*)} = min{*ν*(*S*), *ν*(*D*)}. This maximum value in the overlapped area occurs if the geometric center of species distribution 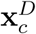 falls in *S_core_ P*(*ν*(*S* ∩ *D*) = min{*ν*(*S*), *ν*(*D*)}) = 1 provided 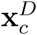 ∈ *S_core_* (Fig. 1). As the assumption of random placement of species distribution center, the probability that the overlapped area is its maximum value is proportional to the area of the *S_core_* in *S_out_*. Therefore, we have the following relationships:

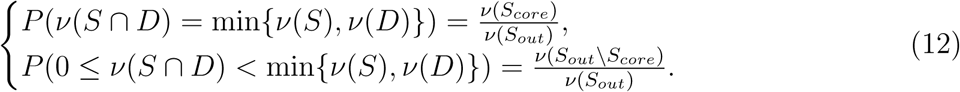

**Figure 2:**
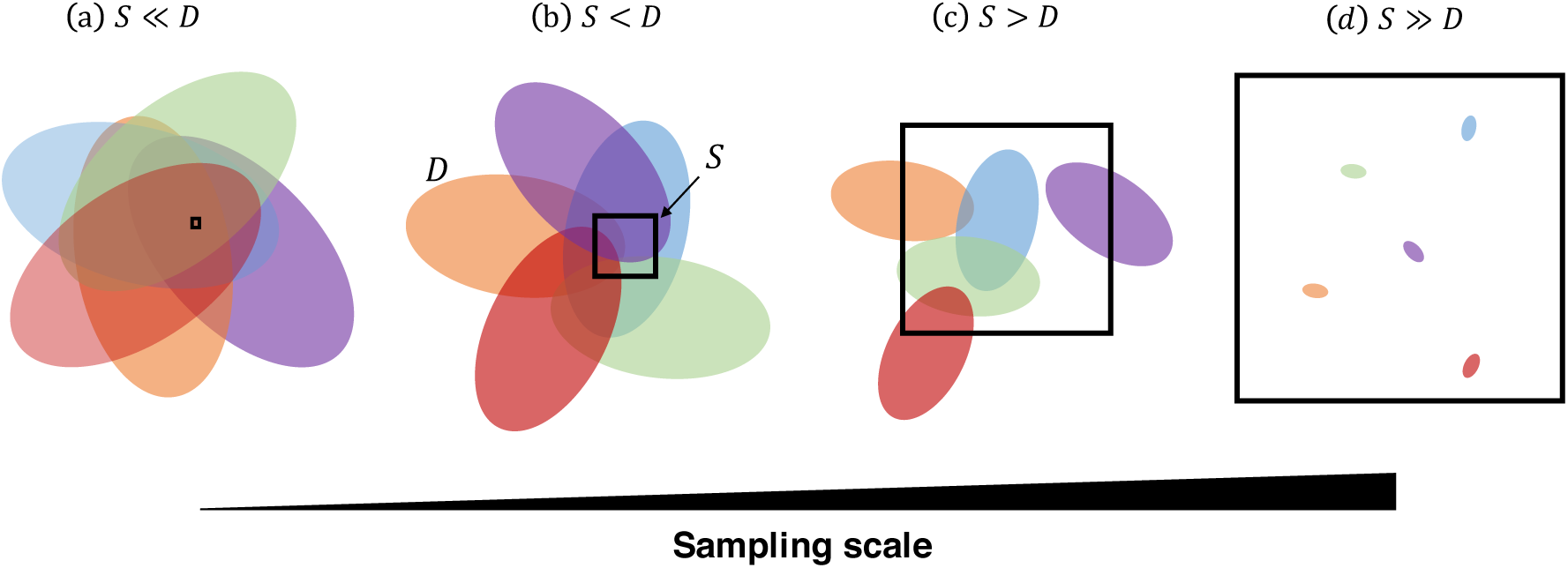
Schematic representation of typical geometric pattern in the derivation of the relative species abundance across sampling scales. The sampling region *S* and distribution range *D* are described by square and ellipse as an example. In each panel, different sampling scales are shown, but the size of distribution ranges remain the same. (a) and (d): when the sampling scale is sufficiently small or large, all sampled species have the same overlapped distribution range with the sampling region (*ν*(*S*) and *ν*(*D*), respectively). (b) and (c): in intermediate sampling scales, this overlapped region is different between sampled species.

As implied in Fig. 2a and d, when *S* ≪ *D* or *S* ≫ *D* is satisfied, *P*(*ν*(*S* ∩ *D*) = min{*ν*(*S*),*ν*(*D*)}) ≃ 1 is satisfied. The abundance of a species within overlapped area *ν*(*S* ∩ *D*) shows variation, and its pdf corresponds to the relative species abundance in area *ν*(*S* ∩ *D*). Let *P*(*x* | *ν*(*S* ∩ *D*)) be the probability of finding a species with *x* individuals in the sampling region *S* given the overlapped area *ν*(*S* ∩ *D*). Then the RSA provided an arbitrary sampling area *ν*(*S*) is described as

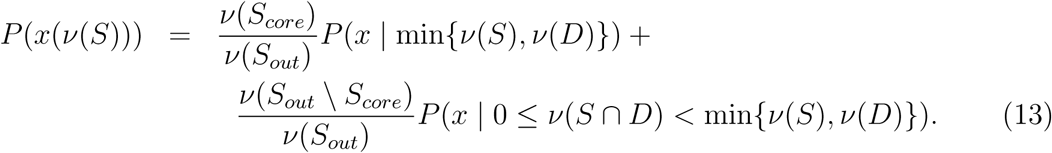

Especially, when *S* ≪ *D* or *S* ≫ *D* is satisfied, Eq. (13) is simplified as

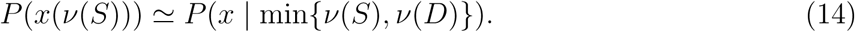

In line with the discussion above, it is straightforward to incorporate the species variations such as the distribution range *D* and biological parameters **b**. By introducing pdfs these parameters follow: *p*(*D*) and *p*(**b**), assuming that the stochastic variable *x* is now a function of **b**, *x*(**b**), and averaging Eq. (13) by the pdfs, we obtain

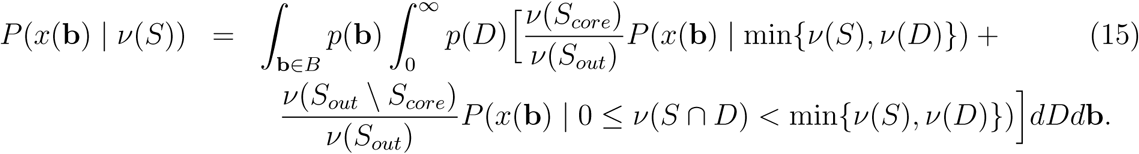

## 3 Generating species distribution patters

To examine the general theory of the geometric approach developed above, we need to introduce a point field *f* (x) defined above that holds information of individual distributions of all species. For this purpose, we make use of point processes [23, 24], a set of spatially explicit stochastic models that generate various point distribution patterns such as random and clustering patterns. One of the advantages of this models is that point processes are amenable to mathematical analysis and easy to implement numerical simulations. In addition, there are a number of applications to ecological studies [11, 19, 20, 22, 25], and models can provide consistent patterns with observed SARs or a population occupancy probability regardless of simple assumptions [11, 22, 25].

Here, we examine the theory developed using two individual distribution patterns: random and clustering distribution patterns, described by the homogeneous Poisson process and the Thomas process, respectively. The former is often used to develop a simplest possible model (e.g., [26]), or to see how the simple assumption deviates from more biologically reliable models (e.g., [16, 19, 22]). As we will see below, deviations of these two distributions in the SAR and RSA are relatively small to make qualitative discussions and these appear in small scales.

The homogeneous Poisson process and Thomas process are defined using the intensity and intensity measure of species *i*, defined above (Eqs. 1 and 2). Also, as mentioned above, individuals of species *i* is restricted within its distribution range *D_i_.* For example, if species *i* is distributed randomly in region *A*, the probability to find *x* individuals within the region with area *ν*(*A*) follows the Poisson distribution with average *μ_i_*(*A*) = λ_*i*_*ν*(*A*):

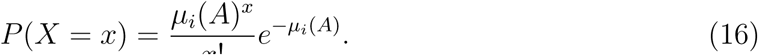

This process generates the homogeneous Poisson process.

On the other hand, the Thomas process is described by the following three steps:

1. Parents of species *i* are randomly placed according to the homogeneous Poisson process with intensity 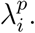
2. Each parent of species *i* produces a random discrete number *c_i_* of daughters, realized independently and identically.
3. The daughters are scattered around their parents independently with an isotropic bivariate Gaussian distribution with the variance 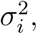 and all the parents are removed in the realized point pattern.

The intensity of individuals of species *i* for the Thomas process is [24]

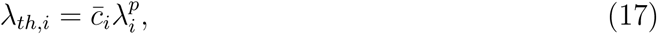

where, *c_i_* is the average number of daughters per parent. To guarantee a consistent number of total expected individuals given area between the two processes, we set 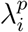 and *c_i_* as

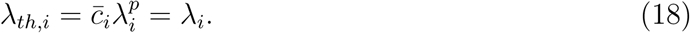

We also assume that the number of daughters per parents *c_i_* follows the Poisson distribution with the average number *C_i_*.

By superposing distributions of all species generated either by the homogeneous Poisson process or the Thomas process, we obtain individual distributions of all species in the whole ecosystem. With these specific individual distribution patterns, we can examine the SAR, EAR, and RSA across scales.

## 4 Emergent macroecological patterns of geometric model

By using the general theory developed, we discuss some specific situations where sampling region and the species distribution ranges are described by circles (Fig. 3a). Also, individual distributions therein are described the homogeneous Poisson process or Thomas process, showing random and clustering distribution patterns, respectively. The assumption of the shapes makes mathematical analysis rather simple and transparent as we can omit the effect of rotation of the species distribution ranges. As we will see below, the second term of the EAR (Eqs. 9, 11) disappears under this assumption. For convenience, all the analyses below are conducted in the polar coordinate. The schematic diagrams under this situation corresponding to Fig. 1 are shown in Fig. 3.

**Figure 3:**
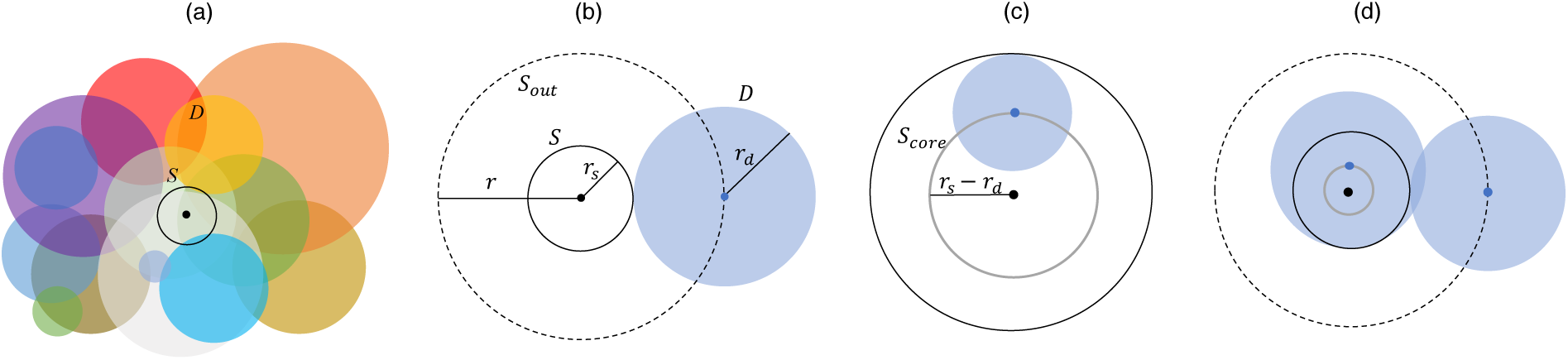
Schematic representations of (a) the sampling region and distribution ranges, and typical geometric relationships used in derivations of the (b) SAR, (c) EAR, and (d) RSA. *S* is the sampling region with radius *r_s_*, *D* is the distribution range of a species with radius *r_d_*. The distance between the centers of *S* and *D* is represented by *r* (in this example, *r* = *r_s_* + *r_d_*). See Fig. 1 for more explanations.

Let the intensity measure of species, *μ_s_*, be a function *f*(*r*,*θ*) defined in the polar coordinate and we assume the homogeneous environment, then the intensity measure of species in the circle with a radius *R* is

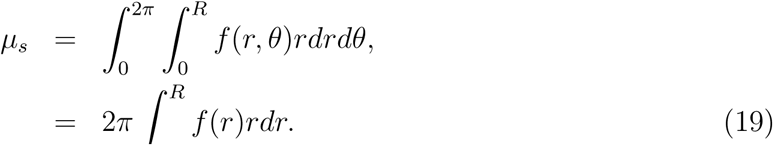

If we define *μ_s_* := λ_*s*_*πR*^2^, where λ_*s*_ is the intensity of species, *f*(*r*) is calculated as

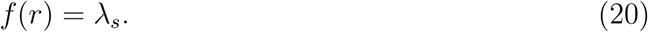

### 4.1 Species area relationship

Let us consider the situation where each species has its distribution area 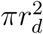 with a radius *r_d_*. We assume that centers of distribution ranges of species are randomly distributed across space, and individual distributions therein show either random or clustering pattern. Then our sampling regime is introduced in such a way that we randomly place the sampling region with radius *r_s_* and count the number of species across spatial scaless by changing the size of sampling region. By doing so, we obtain the SAR (Fig. 3a). From Eqs (19, 20) and with the aid of Fig. 3b, we calculate the number of species *N_s_* found within a sampling unit with area 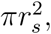 as

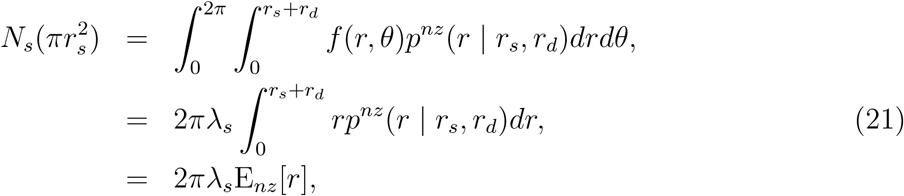

where, *p^nz^*(*r* | *r_s_*,*r_d_*) is the non-zero probability, provided *r_s_* and *r_d_*, that at least one individual of a specie is found given an *r*-distance separation of the centers of sampling and distribution range. Note that 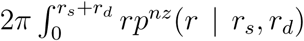*dr* in the second line gives the area corresponding to *ν^nz^*(*S_out_*) in Eq. (5). Eq. (21) implies that the number of species within the sampling region *S* is proportional to the average distance that at least one individuals are found.

We can generalize Eq. (21) to a situation where each species has a different distribution range, therefore different *r_d_*, and biological parameter **b**, (**b** ∈ *B*). Then *r_d_* and **b** follow probability distribution functions, *p*(*r_d_*) and *p*(**b**), respectively. In that situation, Eq. (21) becomes

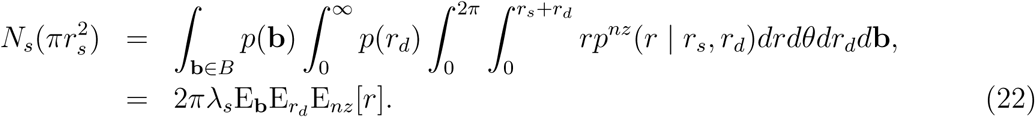

Since this is a special case of the general theory developed above, the discussion about asymptotic slope (Eq. 7) holds.

Here, to show some numerical results in the situation where the distribution range and/or biological parameters differ between species, we examine a situation where the radius of distribution range *r_d_* follows the exponential distribution with the parameter λ_*e*_

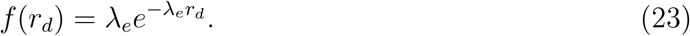

For the biological parameter, we assume that the intensity of each species λ*_i_* follows the gamma distribution:

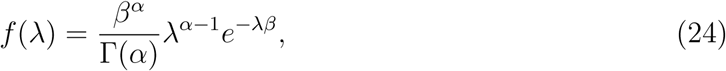

where, *α* and *β* are shape and rate parameters of the gamma distribution respectively. We replace λ by λ*_i_* or λ_*th*,*i*_ depending on the underlying individual distribution process. In addition, we define *c_i_* and 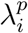 as 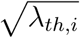 to satisfy 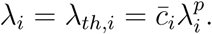 Fig. 4 shows numerically calculated values of neutral (Eq. 21) and non-neutral (Eq. 22) situations, showing the triphasic feature on a log-log plot with different distribution ranges and individual distribution patterns. The third phase of the SAR appears around the average distribution area of species. As discussed above, the slope of each curve asymptotically approaches 1 at large scales. Also, if the distribution range is smaller, it approaches the asymptotic value faster as discussed above. Moreover, it shows that the difference between two individual distribution patterns are relatively small and slight deviations appear only on small sampling scales. Difference between the neutral situation and the non-neutral situation but with only the intensity λ*_i_* are negligibly small when individual distributions is described by the homogeneous Poisson process with the radius of the distribution range *r_d_* = 10km. We checked this holds true for the other curves, and also for different parameter sets of gamma distribution.

**Figure 4:**
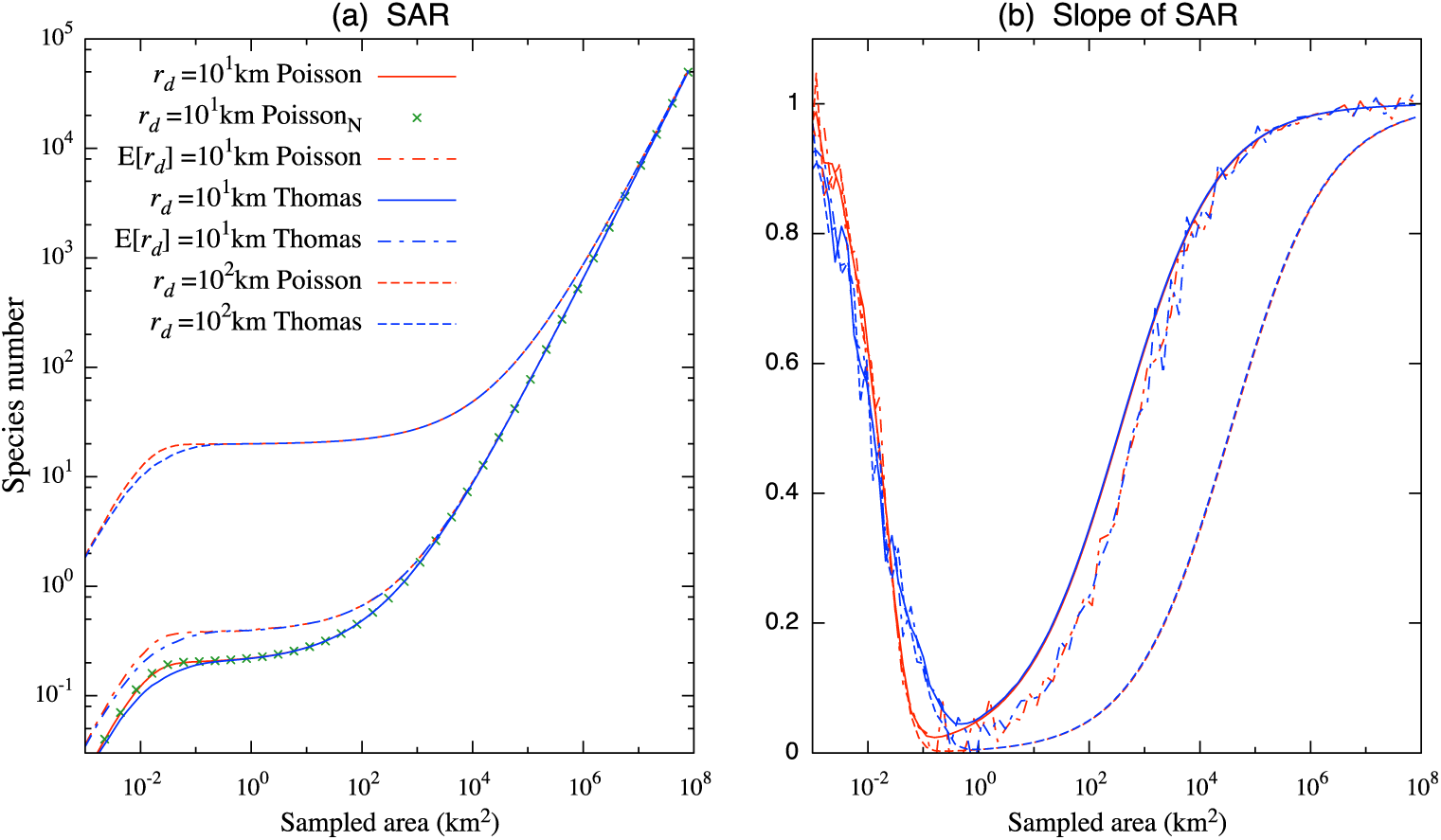
Species area relationship [SAR; (a)] and its slope (**b**) under the homogeneous Poisson process (red and green) and Thomas process (blue), obtained using Eqs. (21) and (22). The intensity λ_*i*_ varies between species according to the gamma distribution except for the neutral situation (star; labeled Poisson_N_). The lines labeled E[*r_d_*] represents the situation where distribution ranges follows the exponential distribution. Otherwise, the distribution range is identical between species (line; *r_d_*=10km line, and dashed; *r_d_*=100km). For the Thomas process, the parameters are λ_*s*_ = 0.00064 (50000 species/π × 5.0 × 10^8^km^2^), λ_*th*,*i*_ = 100, *c_i_* = 10, 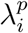 = 10, *σ_p_* = 0.1. For the gamma distribution, we used *α* =10 and *β* = 0.1 (E[λ_*i*_] = 100), and for the exponential distribution λ_*e*_=0.1 (E[*r_d_*]=10) and 0.01 (E[*r_d_*]=10) are used.

### 4.2 Endemic area relationship

When both the sapling region and endemic region are circle, the distribution range of an endemic species satisfies the condition shown in Fig. 3c: the sampling region must be larger than distribution range (i.e., *r_s_* > *r_d_*), and the distance between *S* and *D* must be small enough (i.e., *r* ≤ *r_s_* – *r_d_*). Therefore, the number of endemic species *N_es_* in area 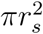 is described as

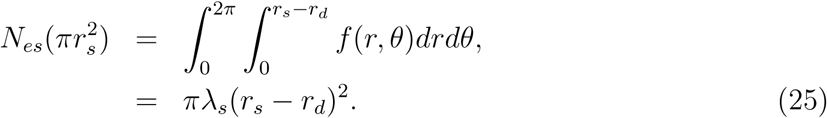

By applying Eq. (6) to calculate the slope of the EAR, it is easily shown provided *r_s_* > *r_d_*

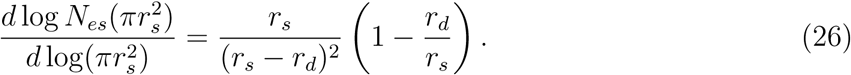

As discussed above, it approach 1 as the sampling region becomes significantly larger than the distribution range: *r_s_* ≫ *r_d_.*

As above, we can generalize the EAR to incorporate variations in distribution ranges of species by introducing a probability distributions, *p*(*r_d_*). However, as noted above, biological parameters does not affect the EAR. For the sake of comparison with the case of the constant *r_d_*, let us assume the average value satisfies E[*r_d_*] = *r_d_.* Eq. (25) now becomes

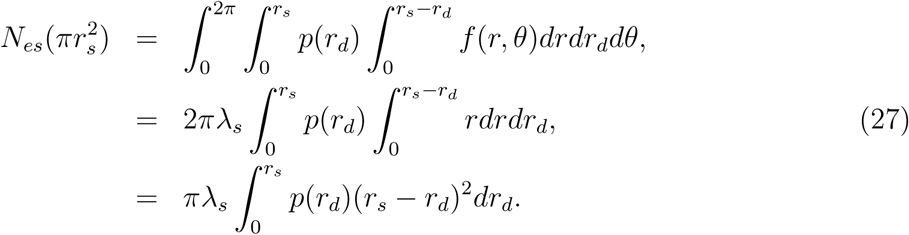

As an example, if we assume the distribution of *r_d_* follows the exponential distribution *p*(*r_d_*) = λ_*e*_*e*^−λ_*e*_*r*_*d*_^, the number of endemic species is

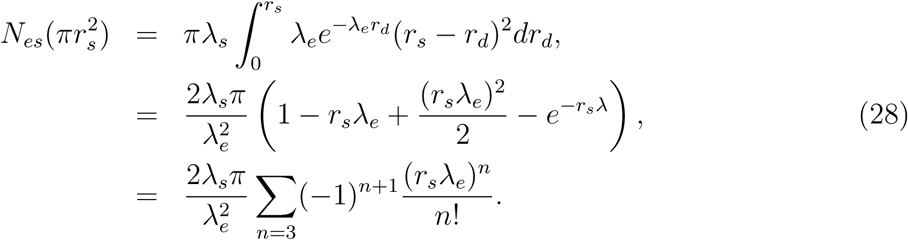

Since 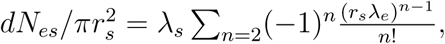 we calculate the slope as

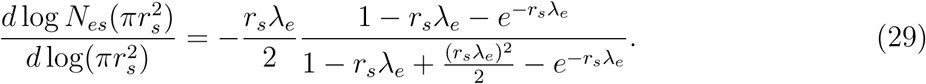

Fig. 5 shows the EAR calculated by Eqs. (25) and (27), and its slope defined by Eqs. (26) and (29). As discussed in General Theory, the EAR on a log-log plot shows asymptotic slope 1, equivalent to the SAR when the sampling region is significantly larger than the distribution ranges *S* ≫ *D.* By the same discussion above, it occurs earlier when the distribution ranges are small. The slope of the EAR becomes infinitely large as soon as the sampling size becomes equivalent to single-sized distribution range. On the other hand, if the distribution range follows the exponential distribution, the slope approaches 1 from a finite value (Fig. 5b), and it also shows that, at small scales, the endemic species is proportional to the sampling area.

**Figure 5:**
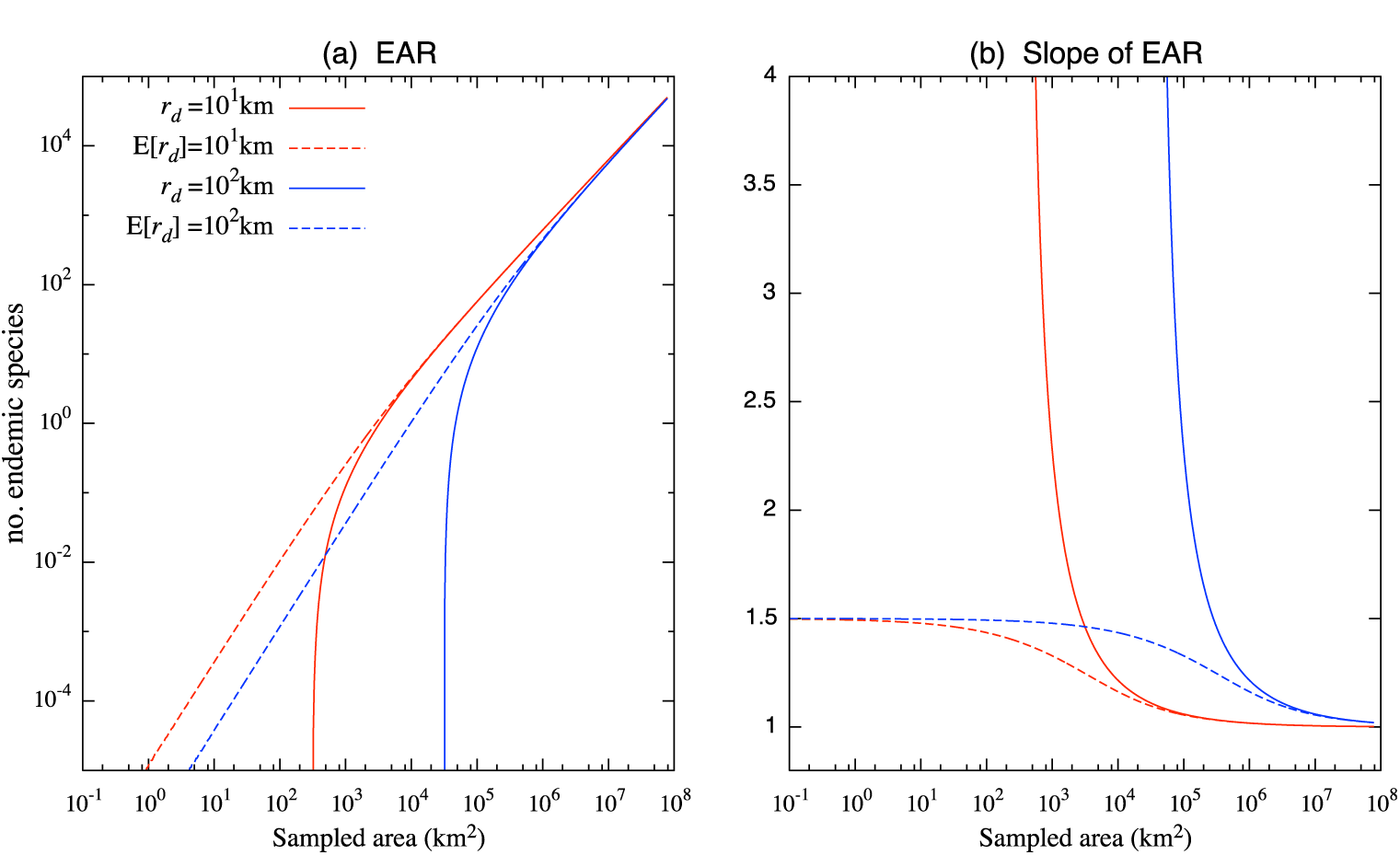
(a) Endemic area relationship (EAR) and (**b**) its slope. Color corresponds to distribution range size (red; *r_d_* = 10km and blue; *r_d_* = 100km). Solid lines represent the situation where no variation in distribution range occurs (Eq. 25). Dotted lines correspond to the situation where the distribution range varies between species, and it follows the exponential distribution *p*(*r_d_*) ∼ exp(–λ*_e_r_d_*) (Eq. 28). To obtain the average value E[*r_d_*] = {10,100}, we set λ_*e*_ = {0.1, 0.01}. For other parameters, the same values are used as in Fig. 4.

### 4.3 Relative species abundance

To derive RSAs across sampling scales, we extensively use mixed probability distributions, especially mixed Poisson distributions [27]. The mixed Poisson distribution was first introduced in ecological study by Fisher et al. [2] in which the celebrated Fisher’s logseries was derived. In our geometric approach, mixed Poisson distributions appear by the nature of multiple stochasticity to determine the number of individuals observed. Namely, there exist intra-species (variations from the expected number of individuals) and inter-species variations (variations of the expected number of individuals itself) of individuals within an overlapped area between the sampling and distribution region *ν*(*S* ∩ *D*). In addition, the overlapped area also varies between species, and the probability distribution of the area *P*(*ν*(*S* ∩ *D*)) depends on sampling scale as shown in Fig 2. Hence, to we need to resolve the scale effect to derive the RSA across scales.

Mixed Poisson distributions are known to produce a variety of pdfs [27], and it is hard to make an exhaustive list of potential RSAs. However, we can consistently obtain RSAs in arbitral sampling scales *S* by scaling up/down, once all the parameters are provided. Therefore, we will firstly focus to recover some well documented RSAs, when the sampling region is much smaller than the distribution range *S* ≪ *D*, probably the most common situation in practice. And secondly, we will scale up the sampling region *S*, by keeping the same assumption, to see how RSA will change. In the first step, we will recover the negative binomial distribution [28], Fisher’s logseries [2], and Poisson lognormal distribution [29].

In the following analysis, we assume that individual distributions are random: it is described by the homogeneous Poisson process. However, as we will see below, the difference of RSAs between the homogeneous Poisson process and Thomas process are small to discuss qualitative features of the RSA, except for the small scales where the deviations of two processes appear in the SAR (Fig. 4).

#### 4.3.1 Some basic properties of mixed Poisson distribution

Here, we introduce some basic properties of mixed Poisson distributions used in the following analysis. For a mixed Poisson distribution, the Poisson intensity itself also follows a probability distribution *g*(λ), and the probability is described [27]

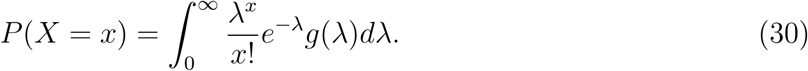

As in [27], we denote this mixture as

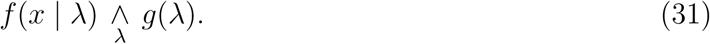

Note that in our application of the homogeneous Poisson process where the probability variable is the number of individuals, the intensity λ in Eq. (30) is replaced by the intensity measure *μ_i_* = λ_*i*_*ν*(*A*) where λ*_i_* and/or *ν*(*A*) vary independently. To specify the probability variable, we use the following expressions:

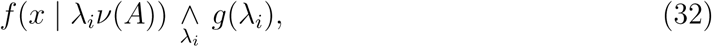

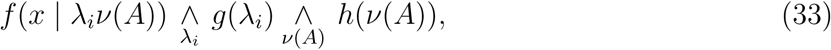

where, the first expression is the case for only λ*_i_* varies and the second expression is the case for both λ*_i_* and *ν*(*A*) vary.

From the Proposition 1 and 2 in Appendix, the probabilities of the mixed Poisson distribution shown in Eqs. (32) and (33) are described

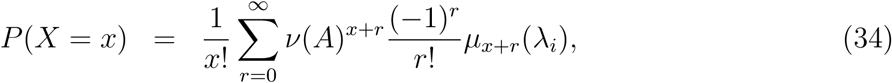

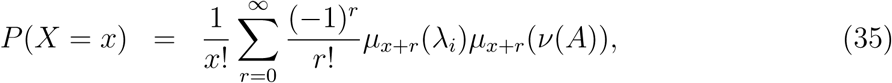

respectively, where *μ_r_*(*x*) is the *r*th moment of *x* about the origin. As we see below, *ν*(*A*) = min{*ν*(*S*), *ν*(*D*)} in our application, and these are RSAs we explore.

#### 4.3.2 Sampling region is much smaller than distribution range: *S* ≪ *D*

In our framework, if species *i* is sampled, the distribution range of species *i* must be overlapped with the sampling region *P*(*S* ∩ *D_i_* = *ϕ*) ≠ 0. Provided by *P*(*S* ∩ *D_i_*, = *ϕ*) ≠ 0, and if the sampling region is much smaller than the distribution range, the sampling region is completely covered by the distribution range *P*(*ν*(*S* ∩ *D_i_*) = *ν*(*S*)) = 1 as in Fig. 2a. In that situation, we can neglect the variations of the overlapped region between species, and consider only the abundance variations of inter- and intra-species within the the sampling region *S*. Namely, this is the situation described by Eq. (32) and the RSA under this condision is obtained by calculating Eq. (34).

##### Negative binomial (Poisson-gamma) distribution and Fisher’s logseries

It is well known that the mixing the Poisson distribution with the gamma distribution produces the negative binomial distribution [27]. By taking certain limits of the negative binomial distribution, Fisher et al. [2] obtained the Fisher’s logseries. Here, we follow the same idea to derive these two distributions. Firstly, by Eq. (30), mixing the Poisson distribution with the gamma distribution Eq. (24) gives the following probability function

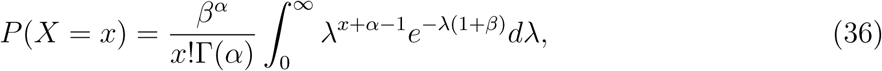

In our situation, the mixture is described by Eq. (32), not by Eq. (31), and the probability of finding a species with *x* individuals are described

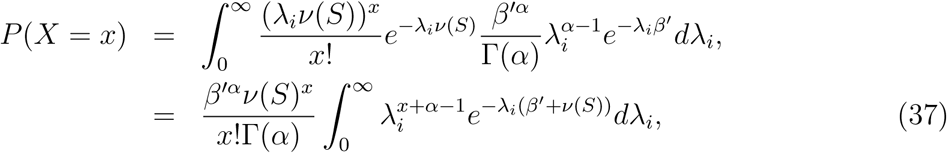

where, we set *β*′ = *βν*(*S*) and changing the variable as *ν*(*S*)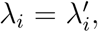 we recover the form of Eq. (36). Eq. (36) is well known to produce the negative binomial distribution

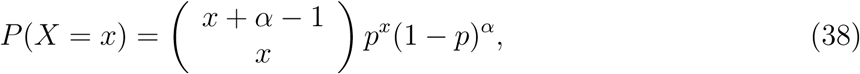

where, in our case *p* = *ν*(*S*)/(*ν*(*S*) + *β*).

To obtain Fisher’s logseries from Eq. (38), we need to further assume the sampling effect: the number *s* of individuals are sampled. By taking the Fisher’s limit [30], we obtain:

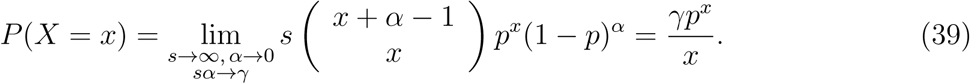

##### Poisson-lognormal distribution

Poisson-lognormal distribution is obtained by mixing the Poisson distribution with the lognormal distribution 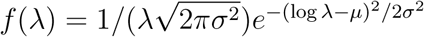[29];

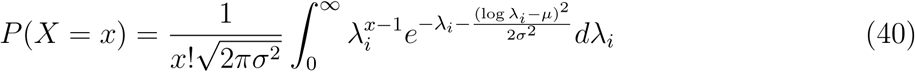

In our situation, the mixture is as in Eq. (32), and it is described

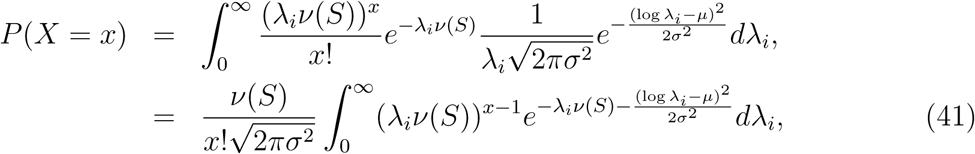

where, by setting *μ* = *μ*′ – log*ν*(*S*) and 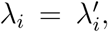 we recover the form of Eq. (40). Therefore, if the parameter λ*_i_* follows the liognormal distribution and sampling region is much smaller than the distribution range, we expect to observe an RSA curve that follows the Poisson-lognormal distribution.

#### 4.3.3 General situation

With the result of the analysis above, we explore more general situations where the sampling region does not necessarily satisfy *S* ≪ *D.* In that situation, the probability function of finding a species with *x* individuals has the form of Eq. (13), in which as in Eq. (14), the above-mentioned situation *S* ≪ *D* (and *S* ≫ *D*) is treated as a special situation. The weighting terms Eq. (12) in this example is easily calculated with the aid of Fig. 3:

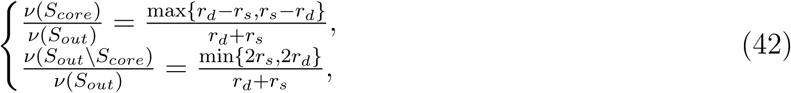

where, one can easily see that the second weighting term disappears when *S* ≪ *D* and *S* ≫ *D*, namely *r_s_* ≪ *r_d_* and *r_s_* ≫ *r_d_*, respectively. The first probability function of Eq. (13) corresponds to the situation where we can neglect the variation of overlapped region, and this situation was already discussed above. Namely, it is described by Eq. (34). The second probability of Eq. (13) corresponds to the situation where the overlapped area varies between species. As we discussed above, the mixed probability distribution has the relationship of Eq. (33), and the probability of finding a species with *x* individuals is described by Eq. (35).

For simplicity, we focus here on the case where the λ_*i*_ follows the gamma distribution as in the first example above (Poisson-gamma), since this case is mathematically more amenable than the case of Poisson-lognormal, and therefore it allows us to discuss results in a mathematically more transparent way. This sole assumption, however, produces a variety of RSAs including similar forms to the negatively skewed lognormal distribution [6], or a left-skewed bell shape curve on a logarithmic scale, resembling the Poisson-lognormal distribution.

To make use of Eq. (35), we need the *r*th moment of a probability distribution of the overlapped area *ν*(*S* ∩ *D*). Fig. 6a shows an example of the relationship of the overlapped area *ν*(*S* ∩ *D*) given an *r*-distance separation of geometric centers between the sampling region and the distribution range of species. Since *r* is randomly given provided that *ν*(*S* ∩ *D*) ≠ *ϕ*, *ν*(*S* ∩ *D*) is a probability variable. We can therefore treat the graph (Fig. 6b), obtained by rotating Fig. 6a and normalizing to 1, as a cumulative distribution function of *P*(*ν*(*S* ∩ *D*), and infer the pdf. Especially, since this function has two different phases, we decompose it into two parts. The region described with *δ* corresponds to the first probability function in Eq. (13) in which the probability is described by a delta distribution and has a peak at min{*ν*(*S*),*ν*(*D*)}. The rest part corresponds to the second probability function in Eq. (13), and it has a similar shape to an arcsin function. Because we require the moment of a probability distribution function, *P*(*ν*(*S* ∩ *D*), it may be a reasonable to approximate it by an existing probability distribution. Therefore, we apply the arcsin distribution, which is a special case of the beta distribution [31]

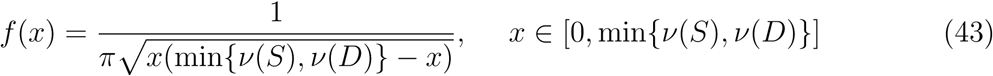

**Figure 6:**
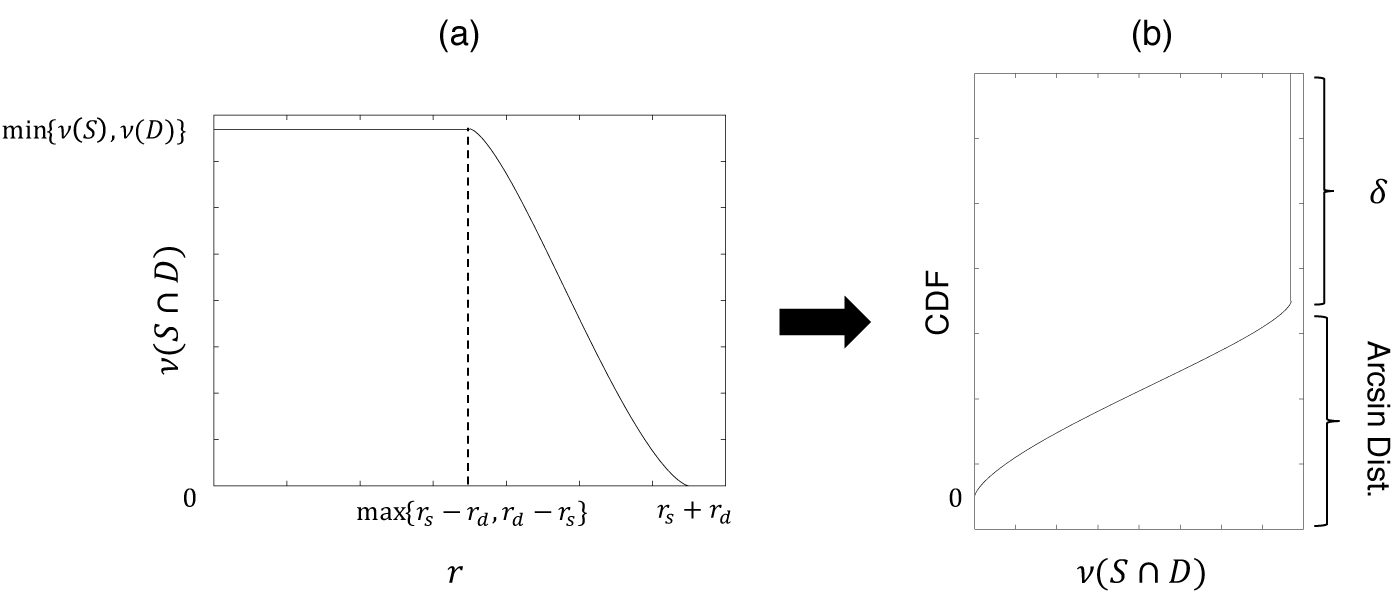
An example form of the overlapped area *ν*(*S* ∩ *D*) given an *r*-distance separation of geometric centers between the sampling region and the distribution range of the species (a), and the cumulative distribution functions of overlapped area (b).

where, the support is defined as *x* ∈ [0,min{*ν*(*S*),*ν*(*D*)}]. The *r*th moment of Eq. (43) about the origin is, by the Proposition 3 in Appendix, described by 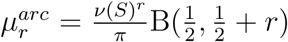 where, 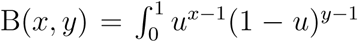*du* is the beta function. By substituting this equation, together with the *r*th moment of the gamma distribution Eq. (24), 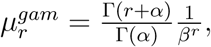 into Eq. (35), we obtain the following probability function (see Appendix for the detailed derivation):

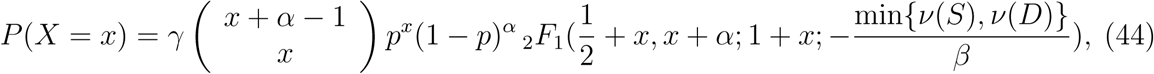

where, *γ* is the coefficient *γ* = 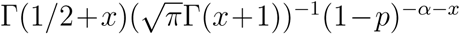 and this part is simplified for *x* ≥ 1 as *γ* = *x*^−1^B(1/2, *x*)^−1^(1–*p*)^−*α*–*x*^, and *p* is now becomes *p* = min{*ν*(*S*), *ν*(*D*)}/(min{*ν*(*S*), *ν*(*D β*). The second to forth factors correspond to the negative binomial distribution the form corresponds to Eq. (38), and _2_*F*_1_ is the Gauss hypergeometric function. We call this distribution the Poisson-gamma-arcsine distribution. From, Eqs. (38), (42), and (44), we derive the general form as in Eq. (13):

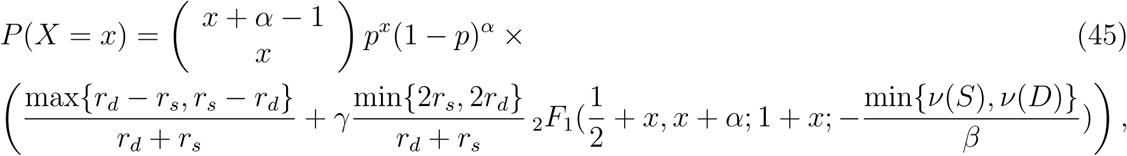

where, the probability distribution is a weighting sum of the negative binomial distribution and the Poisson-gamma-arcsine distribution. As already discussed above, *S* ≪ *D* (*S* ≫ *D*) is the special case of Eq. (45). That is, by taking *r_s_* ≪ *r_d_* (*r_s_* ≫ *r_d_*), Eq. (45) becomes

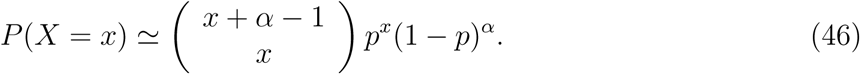

Namely, the RSAs follow the same probability distribution when the area of sampling region and distribution range are significantly different. It is worth emphasizing that to derive Eq.(46), the term approximated by arcsine distribution disappears. Since all the parameters except for *ν*(*S*) are consistent across spatial scales, the difference between these two cases is only in *p* = min{*ν*(*S*),*ν*(*D*)}/(min{*ν*(*S*), *ν*(*D*)} + *β*). It is expected that *p* is close to 1 if *ν*(*D*) = min{*ν*(*S*),*ν*(*D*)}. The same discussion can be made when the intensity follows the Poisson-lognormal distribution Eq. (40), and this is discussed in Appendix (see Poisson-lognormal-arcsine distribtuion), but with mathematically less tractable and computationally intensive form.

Examples of RSAs derived by Eq. (45) across scales (the radius of the sampling region is *r_s_* = 0.05,0.1,0.5,10,100 and 1000km, respectively) are shown in Fig. 7, associated with numerically obtained RSAs provided that the underlying individual distributions are the homogeneous Poisson or Thomas processes. Since numerical calculations with large distribution ranges are computationally expensive, we set *r_d_* = 10km. However, this is suffice to check our analysis. Especially, we examined the zero-truncated form of Eq. (45), since in practice we do not observe the event *X* = 0. To do this, we multiply each distribution by (1 – *P*(*X* = 0))^−1^, where *P*(*X* = 0) is (1 – *p*)*^α^* for Eq. (38) and 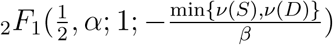 for Eq. (44), respectively.

**Figure 7:**
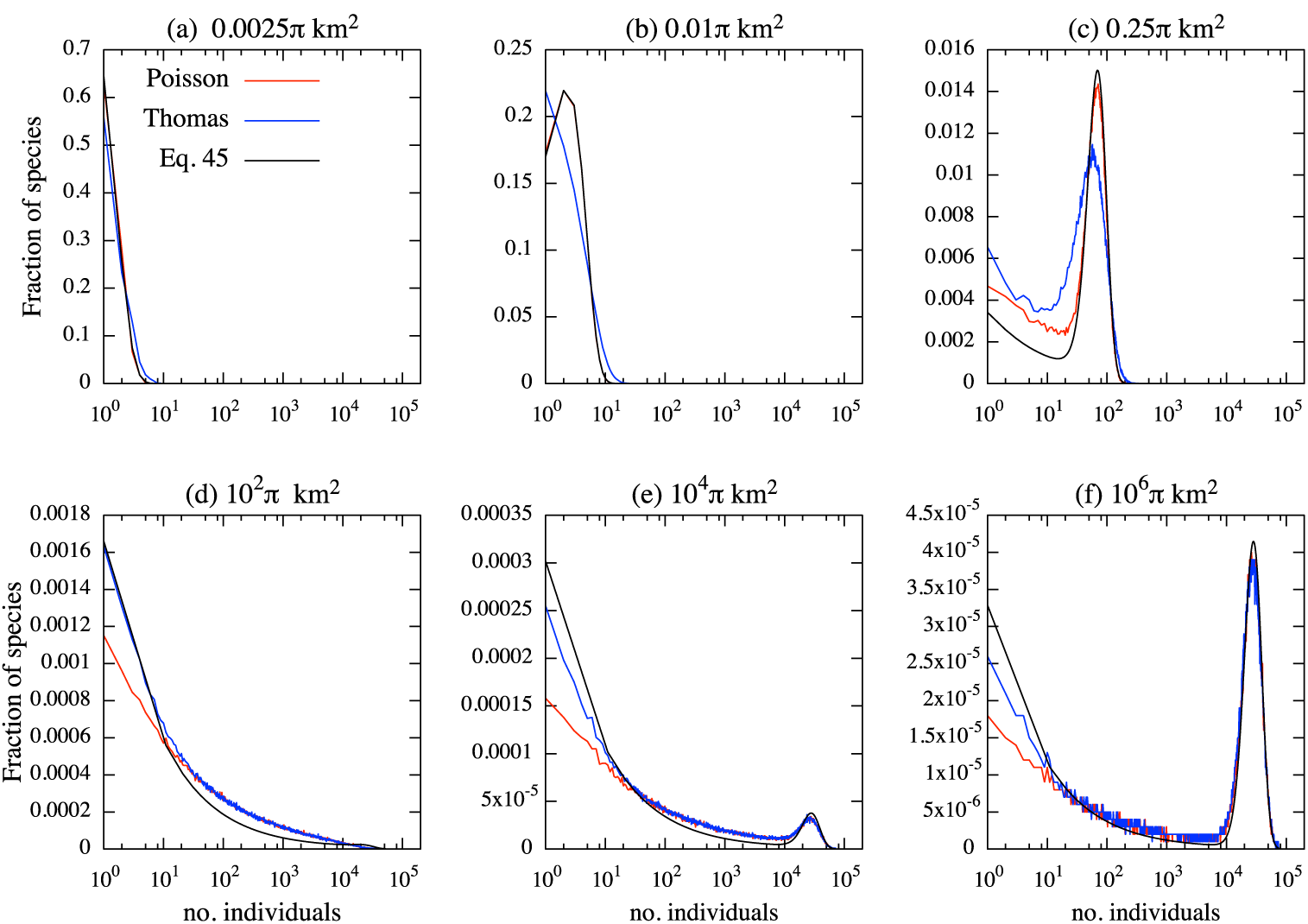
Relative species abundance across sampling regions. The radius of the sampling region in each panel (a)-(f) is *r_s_* = 0.05, 0.1, 0.5,10,100 and 1000km, respectively, and these values are chosen so as to make transitions of the RSA tractable. Each panel shows three different curves regarding to the situations where underlying individual distributions are the homogeneous Poisson process and its theoretical form (Eq. 45), and Thomas process. The theoretical form agrees well with its fully simulated values when the spatial scales of sampling and distribution range are significantly different (i.e., *S* ≪ *D* and *S* ≫ *D*), in which Eq. (45) approaches to Eq. (46) where no approximation is made. Outside these regions, the theoretical curve still captures qualitative aspects of the numerical results, but with different degrees. See the main text for further explanations.

In general, the homogeneous Poisson process and Thomas processes show qualitatively similar curves except for Fig. 7b where deviations between two processes in the SAR appear (Fig. 4). As expected, when the spatial scales of sampling and distribution ranges are significantly different (i.e., Figs. 7a and b; *S* ≪ *D*; the weighting term is (*r_d_* – *r_s_*)/(*r_d_* + *r_s_*) < 0.02) the analytical results show good agreement with numerical results, as Eq. (45) approaches the non-approximated RSA (i.e., Poisson-gamma distribution; Eq. 46). Outside this region, the effect of approximation appears (Figs. 7c-f), showing deviations from the numerical results especially for small *x*, but it still describe qualitative aspects of each RSA. The tail on small *x* (Fig. 7f) disappears in the large limit of the sampling region (Eq. 46) Fig. 8, showing a left-skewed bell shape on a logarithmic scale.

**Figure 8:**
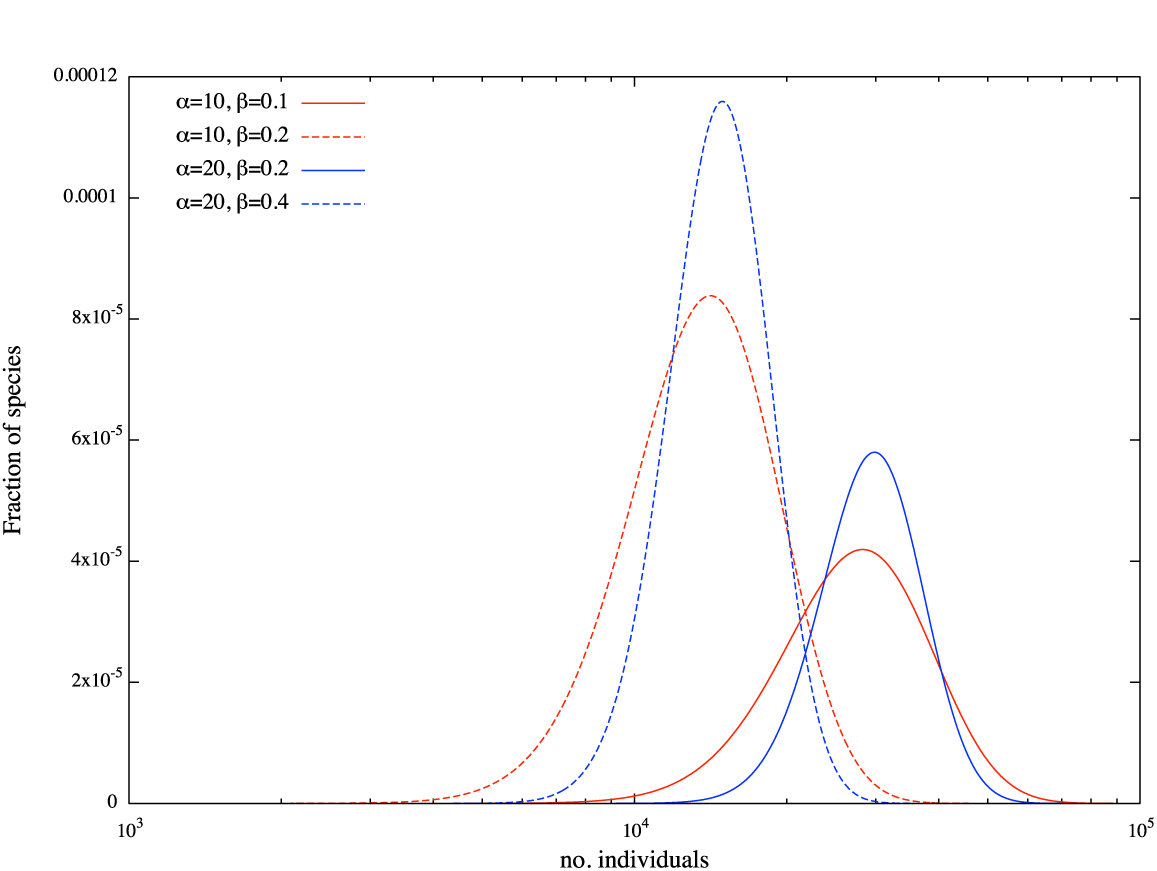
Relative species abundance in the large limit of the sampling region (Eq. 46) with four different parameter sets (*α*, *β*) of the gamma distribution.

## 5 Spatial scaling of *β* diversity: An application of geometric approach

*β* diversity is an important concept in community ecology and conservation biology that describes variations of species compositions between multiple assemblies across spatial scales [13, 32]. The spatial variations inherently include the scaling effect of the size of concerned region and its subregions, and its effect and relationship between other macroecological properties is often of interest of community ecologists [13]. However, data is often not sufficiently enough to unveil the relationship empirically, and as far as we know, there is no theoretical attempt to explore its property (but Barton et al. [13] provided conceptual discussions). Here, to demonstrate potential uses of the geometric model developed, we apply the model to discuss this issue. For this purpose, our analysis is restricted to two scenarios: same parameter values are used as the results of the SARs E[*r_d_*] = 10^1^km^2^ Poisson/Thomas shown in Fig. 4. In the following, we briefly summarize scaling issues of *β* diversity and the diversity index used in the analysis.

### 5.1 Spatial grain and spatial extent

To quantify *β* diversity it requires us to define two spatial scales; *spatial extent* and *spatial grain* [13] Fig. 9. Spatial extent is the scope of our observation, and spatial grain is the unit of sampling within the extent. Once we define the spatial extent, spatial grain, and point patterns, we obtain the matrix *P*: provided that there are *s* species in the spatial extent, and arbitrarily dividing the community into *n* assemblies, the occurrence probability of species *i* in community *j*, *p_ij_* = 1 (Σ_*i*_*p_ij_* = 1) creates the following matrix

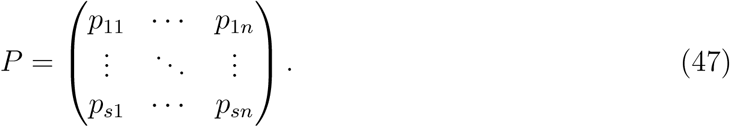

**Figure 9:**
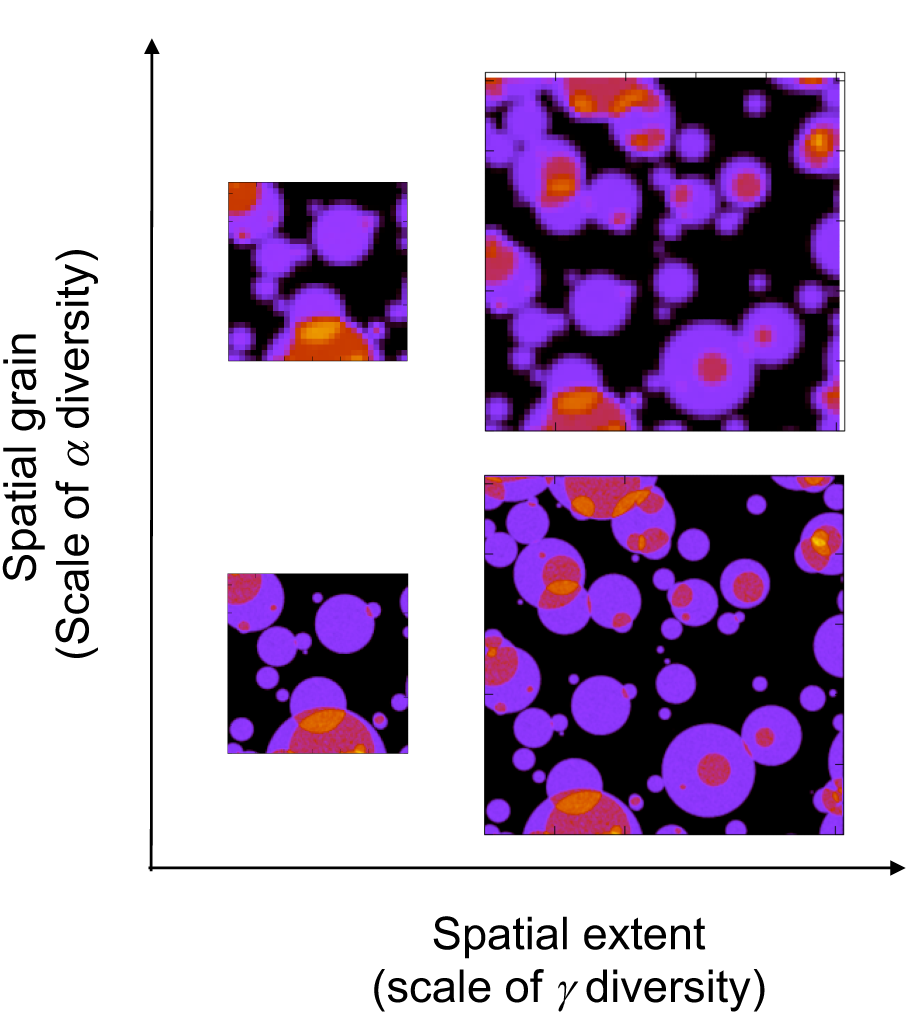
Schematic figure of spatial extent and spatial grain. Colors indicate the abundance of all the species within the spatial extent 128km×128km (left) and 256km×256km (right). Spatial grain is 1km× 1km (bottom) and 4km×4km (top).

Changing the size of spatial extent and/or spatial grain changes the size of the matrix *P*, since a smaller spatial extent may hold a fewer number of species, and a fine spatial grain increases the number of sampling patches. Therefore, if we change either one or both of these scales, the matrix *P* is changed into another matrix *P*′. We describe this operation as *P* ↦ *P*′. For example, if Eq. (47) is the *s* × *n* matrix with equal-sized patches and *n* is an even number, and if its spatial grain is doubled, the operation *P* ↦ *P*′ gives a *s* × *n*/2 matrix.

In practice, each patch has a different significance on a diversity index, and this is described by the weight vector **w** = (*w*_1_,*w*_2_, …, *w_n_*). We can also define the same operation for the weight vector associated with a scale change.

### 5.2 Diversity indices

Statistical discussions over its reliable definitions given a huge number of different definitions [33] and bridge between different definitions has been actively discussed [32, 34-37]. Here we adapt the definition of Jost [32]. Jost [32] showed that when weights in diversity indices are unequal, meaningful diversity index is only the Shannon measure since this only satisfies five requisite conditions for diversity indices. The Shannon measure is described by

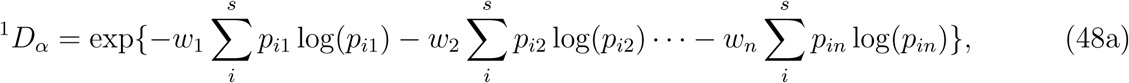

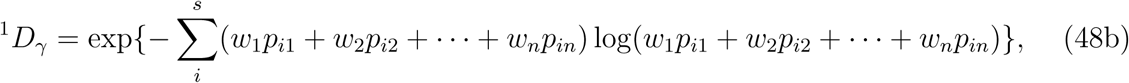

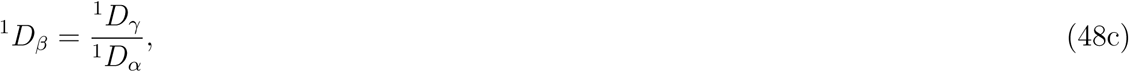

In order to discuss the effect of a scale change of the spatial grain consistently, we further need a condition for independence of a choice of weights on the gamma diversity Eq. (48b). That is to say, scale changes of the spatial grain, provided a consistent spatial extent, should not affect the gamma diversity. It is easy to see this requisite is satisfied if we define the weight vector by population abundance of each patch, *w_j_* = Σ_*i*_ *N_ij_*/ Σ_*i*,*j*_, where *N_ij_* is the abundance of species *i* in patch *j*. Furthermore, this is the only choice to hold this condition (Theorem 1).

Since *β* diversity depends on the number of spatial grains, we require normalize the *β* diversity onto [0,1] to make a meaningful comparison of spatial variations [32]. When all the weighting terms are not identical, the regional homogeneity measure (1/^1^*D_β_* – 1/^1^*D_w_*)/(1 – 1/^1^*D_w_*) [32] may be used for the normalized measure of *β* diversity, where ^l^*D_w_* is the Shannon measure of weighting term ^1^*D_w_* = exp(– Σ*_j_ w_j_* log(*w_j_*)). We use its complement

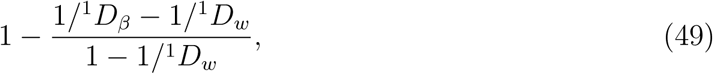

as a relative inhomogeneity measure. This measures is 0 if all the communities are identical and 1 if all the communities are distinct.

### 5.3 Normalized *β* diversity across spatial extent and grains

To see how the normalized *β* diversity (inhomogeneity measure) changes, we compute Eq. (49) across spatial extents and spatial grains. We choose the range of spatial extent to cover three different phases in the SAR (Fig. 4) (2^−6^km^2^ – 2^16^km^2^), and we divided the spatial extent into equal-sized subregion with spatial grain size 2^−8^km^2^ – 2^*s*–2^km^2^, where *s* determines the size of spatial extent, so that each scenario has minimum 4 subregions. To compute inhomogeneity measure across spatial extent and spatial grains, we used only one realization of point patterns by taking the following three steps: (i) define point patterns in the maximum spatial extent (e.g., defined on the plane [0, 2^8^] × [0, 2^8^]); (ii) define a spatial extent (e.g., on the plane [0, 2^−1^] × [0, 2^−1^]); and (iii) calculate the inhomogeneity measure for each spatial grain with area (2^−8^, 2^−6^, 2^−4^). In the step (ii), we started from the minimum spatial extent 2^−8^km^2^, and repeated the step (ii) and (iii) until spatial extent reached the maximum extent 2^16^km^2^. We eliminate the situation where no individual exists when the minimum spatial grain size 2^−8^km^2^ is applied to make sure each scenario contains at least one individual. When all individuals situated in one subregion, we define the inhomogeneity measure as 0.

Fig. 10 shows the numerically calculated inhomogeneity measure averaged over 500 simulation trials, associated with the SAR curves of underlying processes. Overall, the underlying processes under concern does not cause prominent qualitative effect. The top two figures show heat map of the inhomogeneity measure under the (a) Poisson and (**b**) Thomas processes. In both panels, the inhomogeneity measure shows higher value as the spatial extent increases. Under this operation, the effect of applying different spatial grain becomes small, especially the SAR is in the third phase (the bottom two figures with fixed spatial grains 2^−8^km^2^, 2^−6^km^2^). Conversely, the inhomogeneity measure shows small value when spatial extent underlies in the left-side of the second phase of of the SAR, and its spatial grain is large.

**Figure 10:**
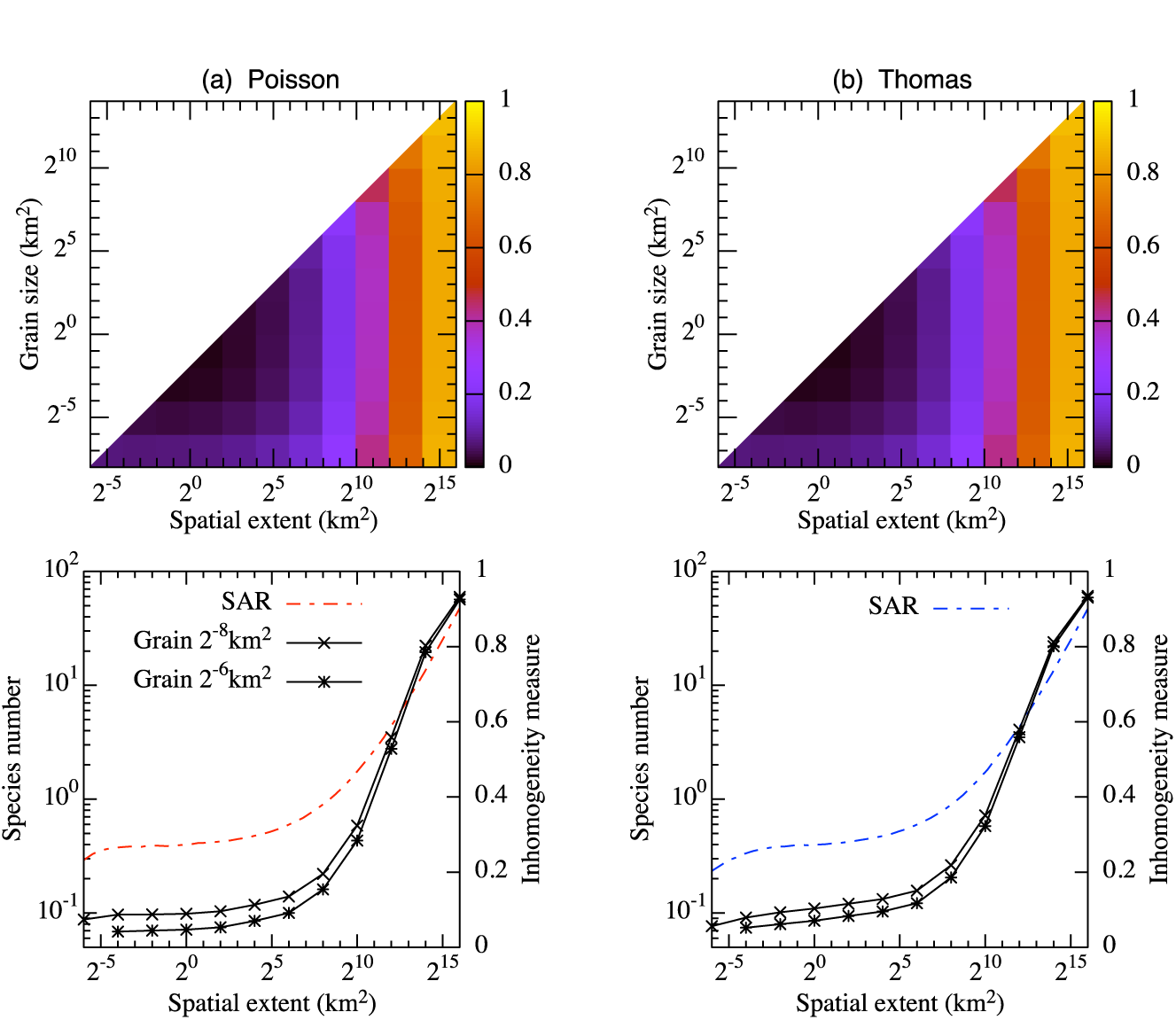
Inhomogeneity measure averaged over 500 simulation trials, associated with the SAR curves of underlying (a) Poisson processes and (**b**) Thomas process. Top two figures show heat map of the inhomogeneity measure across spatial extent and spatial grain. Bottom two figures show some slices of the top figures, where fixed spatial grains (2^−8^km^2^, 2^−6^km^2^) are applied while spatial extent varies. Same parameter values are used as the results of the SARs E[*r_d_*] = 10^1^km^2^ Poisson/Thomas shown in Fig. 4.

## 6 Discussion

We develop a novel framework to derive macroecological patterns across scales by explicitly linking the distribution of individuals within ranges, the size of ranges, and the spatial extent of the sampling region. The model phenomenologically describes species and individual distributions and in the derivations the SAR, EAR, and RSA, no preceding assumptions about these relationships are required. Rather, the model requires a single set of parameters to generate macroecological patterns across scales. Although the model does not explicitly assume specific biological mechanisms such as population or community dynamics, dispersal, and speciation, it still recovers several well-known macroecological patterns including the tri-phasic SAR and Fisher’s logseries [2], the Poisson gamma distribution (negative binomial distribution) [28], Poisson lognormal distribution [29], and similar form to a negatively skewed lognormal distribution [6] as RSAs.

This finding does not imply that the biological mechanisms shaping biodiversity pattern are irrelevant or uninteresting. Rather, our theory demonstrates the minimum assumptions are sufficient to recover such ubiquitous ecological patterns by linking pattern in individuals and species to aggregate macroecological patterns. The latter can be useful as mechanistic theory development as well, as once a mechanistic model has predicted intraspecific point patterns and/or geographic range geometry, our theory can link them to macroecological patterns. That said, it is clear the general forms of commonly-studied macroecological patterns investigated here are general features of biodiversity pattern emerging from the most basic assumptions, and are not indicative of specific ecological processes to the exclusion of others.

We showed that the tri-phasic SAR with and its asymptotic slope on a log-log plot are ubiquitous patterns in the geometric model, emerging all the scenarios examined (Fig. 4a) that encompasses random and cluster individual distribution patterns, and neutral and nonneutral ecological parameters. It can be discussed in line with the findings of Grilli et al. [11] that the tri-phasic SAR is the outcome associated with two bending points in the SAR produced by local and large sampling area that scales the species number provided by a simple geometric realization. Especially, we showed that the third phase in the SAR appears around the average area of the species distribution ranges. This may correspond to the biological interpretations that sampling area exceeds correlation distance of biogeographic process [5, 6]. Notably, we found a negligible difference in the SAR between the neutral situation and a quasi-neutral situation where only the individual intensity (λ*_i_*) varies between species. However, this variation is necessary to derive well-documented RSAs discussed above. This fact suggests that only small effect occurs on non-zero probability that is used in the definition of the SAR (Eq. 8) given underlying ecological assumptions. In addition, we demonstrated that the spatial scaling of beta diversity can be discussed with the tri-phasic features of the SAR curve. Typically the normalized beta diversity (inhomogeneity measure) increases with the spatial extent, and it shows largest value in the third phase of the SAR where the spatial extent exceeds the correlation length of biogeographic process.

In our framework, the EAR is calculated by explicitly taking distribution ranges into account. Under the definition, the EAR on a log-log plot has the same asymptotic slope 1 as the SAR. This result cannot be directly compared to previous studies [10,11,16] where the endemic species is calculated based on the probability to find a species given region where the probability asymptotically approaches 0 as the sampling region approaches 0. On the other hand we apply more straightforward definition: the expected number of species distribution range enclosed completely by the sampling region. Therefore, our definition gives 0 endemic species unless the scale of sampling region exceeds the scale of species distribution range. Asymptotic behavior at small sampling scales can also occur in our definition by assuming a pdf for the species distribution ranges where rather small distribution ranges allow to exist. Although this latter assumption may not be realistic but may be used to discuss about the regional EAR as discussed in He and Hubbell [6]. Difference between the two definitions appear especially in small sampling scales. In fact, Grilli et al. [11] showed that the slope of the EAR converges to the slope of SAR at large scales that agrees with our discussion at large sampling scales.

Importantly, we provide a single equation describing the RSA with an arbitrary sampling scale (Eqs. 13 and 45). The equation is composed of two mixed-probability distributions where their weights are determined by geometries and scales of the sampling region and species distribution range. Specifically, one of the distribution accounts effects of intra- and inter- species variations in an equal-sized region, and the other distribution accounts, on top of these variations, variations of sampled distribution range sizes. In the latter distribution the effect of sampling size is considered: a species that sampled only partially shows smaller abundance than the case that the species is fully sampled. This latter variation causes left-tailed RSAs as some species are sampled only small regions and others are fully sampled (Fig. 7c-f). Once the sampled region becomes sufficiently large enough, the species sampled only partially becomes negligible and the left tail is vanished and it results in lognormal like distribution (Fig. 8) provided each species has sufficiently large abundance. Lognormal or lognormal-like has been claimed as the potential RSA (SAD) at global scales in previous works [38–40]. Specifically, Eq. (45) is the weighted sum of the Poisson-gamma distribution and Poisson-gamma-arcsine distribution. Notably, this single equation does not provide a single pdf corresponding to the RSA but generates multiple pdfs. This property may provide a new insight into a long standing discussion in ecological community about a fundamental RSA pattern of ecosystems, where a number of attempts has tried to fit a single pdf to RSAs across scales (e.g., [8, 28]). In addition, Eqs. (13 and 45) predict that in the two limits of spatial scales *S* ≪ *D* and *S* ≫ *D*, namely the scales of sampling region and distribution ranges are significantly different, the underlying pdfs are equivalent, but with different parameter values. In practice, RSAs in plant communities are typically discussed in relatively small scales (i.e., < 50-ha; e.g. [22, 41]), and this case may be the situation of *S* ≪ *D*, where our results provide Fisher’s logseries, Negative binomial distribution, and Poisson lognormal distribution, depending on the biological assumptions.

Another notable finding is that the prominent variations of different individual distribution patterns (random and cluster) occur only a limited spatial scales in the SAR and RSA. For the SAR, it occurs around the transition points between the first and second phases of the tri-phasic curve (Fig. 4), shows agreement with the results obtained by [22], in which the authors examined the random-placement model and Thomas process to fit an observed SAR patterns up to 50-ha in tropical forests. No, if any, variations between the random and clustered model suggests that there exist spatial scales where some specific biological processes such as the dispersal distance and size of clusters dominate community pattern formations, since these create significantly different geometric patterns within the scope of small scales. However, once the sampling area becomes large enough, e.g., significantly larger than the area dispersal kernel covers, each component of a cluster plays the same role as randomly placed individuals (Note that each parent location placed randomly). Intuitively speaking, if the two geometric patterns generated by the homogeneous Poisson and Thomas process are observed by the scope of such a large scale, it is not easy to say which geometric pattern is which, as long as the number of individuals are the same and the number is large enough to create a certain geometric pattern. This property suggest that random individual distributions can be used to explore the macroecological and community structures with larger spatial scales, and it makes analysis significantly accessible.

In this paper, we developed a theoretical framework to describe macroecological and community patterns across multiple scales with an example for a potential application to a question in community ecology. Other promising applications are for biodiversity conservation, where spatially integrated approaches such as spatial design of reserve networks have been widely discussed [42, 43], and our framework would provide a theoretical exercise of ecosystem management. We minimized the assumptions of the model for its parsimony and keeping analytical tractability. Nonetheless, based on the general theory developed to derive the SAR, EAR, and RSA in an arbitrary situation, our framework can be flexibly extended and examined various ecological assumptions, at least numerically; for example, point processes provide a framework to discuss a heterogeneous environment (e.g., [24]). Rather, one can use any model to generate a point field *f* (x), and examine the macroecological patterns by using the general theory. Such experiments would provide further insights into macroecological patterns and applicability of the model.

## Acknowledgements

We would like to thank Yoh Iwasa for a constructive discussion. NT was supported by Grant-in-Aid for the Japan Society for the Promotion of Science (JSPS) Fellows. Financial support was provided by JSPS (No. 15K14607 to YK and No. 17K15180 to EPE) and Program for Advancing Strategic International Networks to Accelerate the Circulation of Talented Researchers, JSPS (to YK). EPE and NT were additionally supported by subsidy funding to OIST.

## Appendix

Here we describe some propositions which are used in the main text. In the main text, the description *ν*(*A*) used below is replaced by min{*ν*(*S*),*ν*(*D*)}.

### Proposition 1.

Provided that the moments of the mixing distribution in a mixed Poisson model exist, the probability function of the mixture distribution can be written as

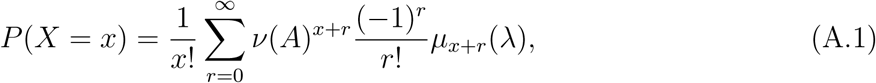

where, *μ_r_*(λ) are the *r*th moment of λ about the origin.

### Proof.

The proof is straightforward from the definition and similar result is found in [27]:

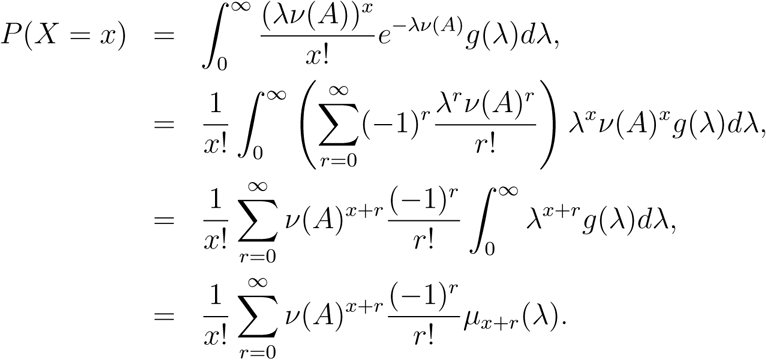

□

### Proposition 2.

Provided that the moments of the mixing distribution in a mixed Poisson model exist, the probability function of the mixture distribution can be written as

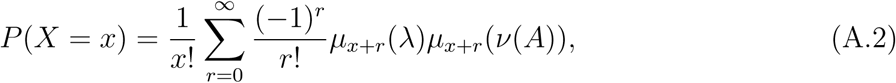

where, *μ*_*x*+*r*_(λ) and *μ*_*x*+*r*_(*ν*(*A*)) are the *r*th moment of λ and *ν*(*A*) about the origin, respectively.

### Proof.

The proof is just an extension of the proposition 1.

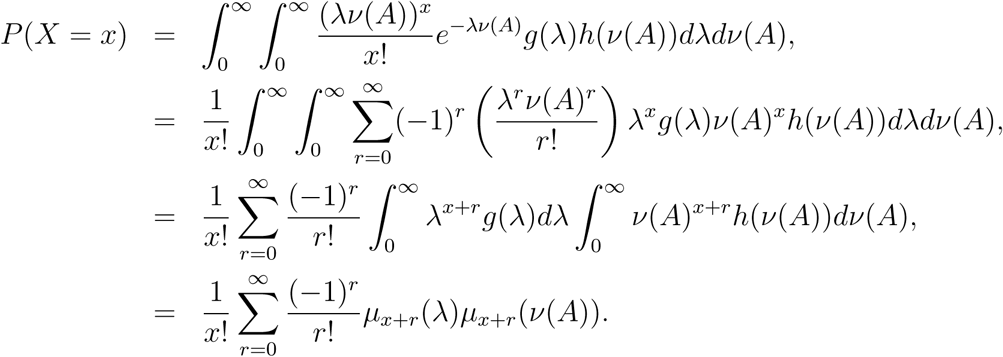

□

### Proposition 3.

The *r*th moment of the arcsine distribution with support *x* ∈ [0,*ν*(*A*)]

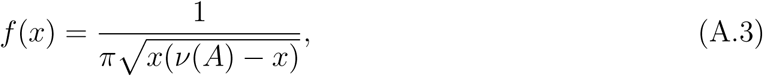

is described by

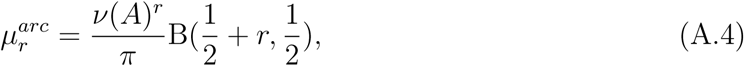

where, 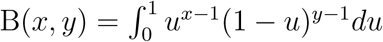 is the beta function.

### Proof.

Using the substitution *w* = *x*/*ν*(*A*),

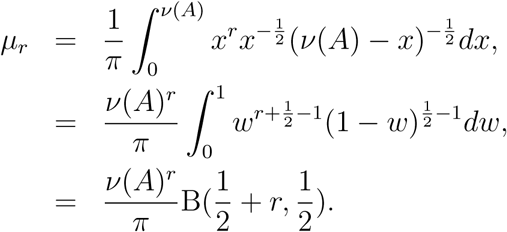

□

### Threorem 1.

*The gamma diversity defined by Eq. (48b) that is independent of the choice of the number of patches or its size is uniquely determined*, *and it is when the weight vector is proportional to the population abundance of each patch.*

### Proof.

Let us assume there exists a vector **w**′ = 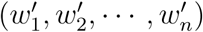 that satisfies 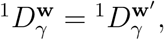 where superscripts **w** and **w**′ indicate weight vector used. By the assumption, we have 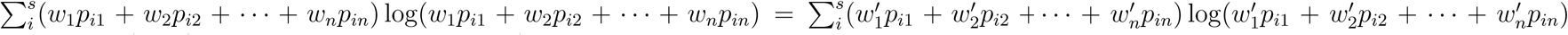 and arranging this expression, it becomes 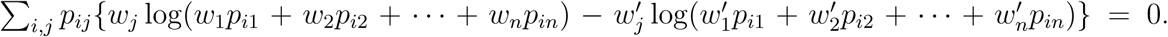 Since *p_ij_* may or may not be zero we must have 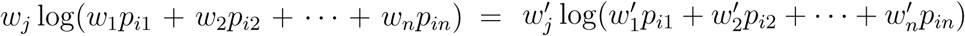 for all *i* and *j*. This is clearly 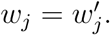
□

### Derivation of the Poisson-gamma-arcsine distribution (Eq. 44)

Now, using Proposition 3 and the *r*th moment of the gamma distribution *f* (λ) = *β^α^*/Γ(*α*)λ^*α*–1^*e*^−λ*β*^

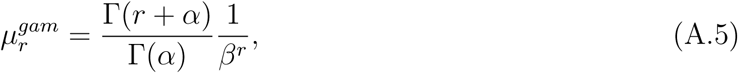

we calculate the probability of mixing distribution Eq. (A.2) with properties of the beta function B(*x*,*y*) = Γ(*x*)Γ(*y*)/Γ(*x* + *y*) and the Gamma function 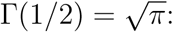

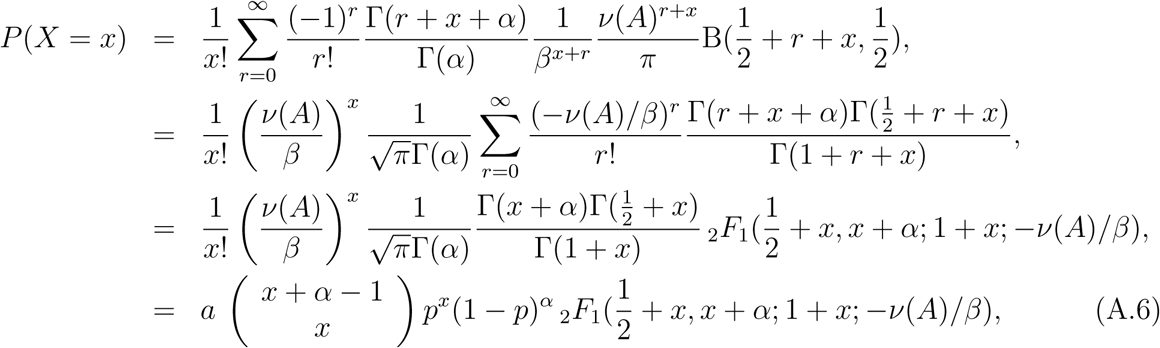

where, *α* is Γ(1/2+*x*)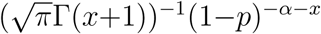 and is simplified for *x* ≥ 1 as *x*^−1^B(1/2,*x*)^−1^(1 – *p*)^−*α*–*x*^, the second to forth factors correspond to the negative binomial distribution with *p* = *ν*(*A*)/(*ν*(*A*) + *β*), and _2_*F*_1_ is the Gauss hypergeometric function. *α* and *β* are the shape and rate parameters of the gamma distribution. Note the negative binomial distribution here is equivalent to Eq. (38).

### Poisson-lognormal-arcsine distribution

By substituting the *r*th moment about the origin of the arcsine distribution Eq. (A.4), and the Poisson-lognormal distribution 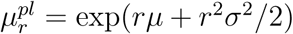 [29] into Eq. (35), we obtain the RSA of an arbitrary form

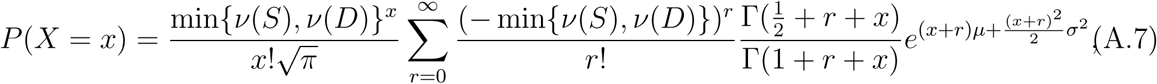

where as in the case of the Poisson-gamma-arcsine distribution, we call this pdf the Poisson-lognormal-arcsine distribution. Using Eqs. (40), (42), and (A.7), we obtain the full form of the RSA across scales

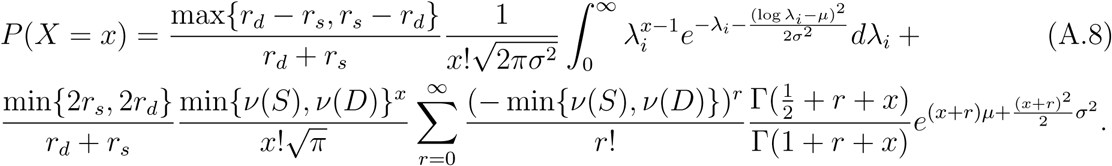

As above, Eq. (A.8) becomes the Poisson-lognormal distribution in the limits of *S* ≪ *D* and *S* ≫ *D*:

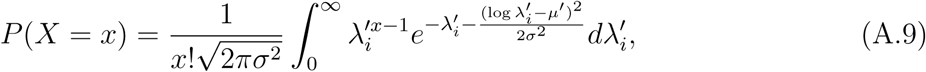

where, by setting *μ* = *μ*′ – log(min{*ν*(*S*),*ν*(*D*)}) and min{*ν*(*S*),*ν*(*D*)}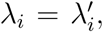 we recover the form of Eq. (40). Therefore, if the parameter λ*_i_* follows the Log-normal distribution and sampling region is much smaller than the distribution range, we expect to observe an RSA curve that follows Poisson-lognormal distribution.

